# Epigenetic constraint of cellular genomes evolutionarily links genetic variation to function

**DOI:** 10.1101/2024.10.28.620690

**Authors:** Enakshi Sinniah, Dalia Mizikovsky, Woo Jun Shim, Chris Siu Yeung Chow, Yassine Souilmi, Fei-Fei Cheng, Zhili Zeng, Jordan Laurie, Matthew Foster, Sonia Shah, Mikael Bodén, Jian Zeng, Bastien Llamas, Nathan J. Palpant

## Abstract

Cellular diversity is a product of evolution acting to drive divergent regulatory programs from a common genome. Here, we use cross-cell-type epigenetic conservation to gain insight into the impact of selective constraints on genome function and phenotypic variation. By comparing chromatin accessibility across hundreds of diverse cell-types, we identify 1.4% of the human genome safeguarded by conserved domains of facultative heterochromatin, which we term regions under “cellular constraint”. We calculate single-base resolution cellular constraint scores and demonstrate robust prediction of functionally important coding and non-coding loci in a cell-type-, trait-, and disease-agnostic manner. Cellular constraint annotation enhances causal variant identification, drug discovery, and clinical diagnostic predictions. Furthermore, cell-constrained sequences share paradoxical evolutionary signals of positive and negative selection, suggesting a dynamic role in driving human adaptation. Overall, this study demonstrates that evolutionary chromatin dynamics can be leveraged to inform the translation of genetic discoveries into effective biological, therapeutic, and clinical outcomes.

## Introduction

Elucidating the functional importance of genomic loci, beyond genetic association to human diseases and traits, is critical to understanding the molecular and cellular basis of phenotypic diversity^1^. While numerous metrics exist to assess the function of protein-coding sequences with high confidence^2,3^, most human genetic variation resides in non-coding regions of the genome with limited interpretability^4^. In recent years, single-nucleotide evolutionary constraint scores have facilitated causal variant discovery and functional interpretation^5–8^ on the basis that variation in conserved sequences is more likely to be deleterious or influence phenotype. However, sequence conservation does not capture all functional elements^9,10^, and requires a large number of eukaryotic genomes to be sequenced to accurately estimate constraint via comparative analysis^11^. Moreover, sequence similarity alone cannot indicate tissue- or cell-type-specific activity. Hence, linking evolutionary constraint metrics to molecular pathways and cell-type-specific function remains a challenge and are underused in cell biology contexts. Comparative epigenomic analyses have emerged as powerful approaches to complement cross-species sequence conservation, offering insights into the evolution of gene regulatory programs at a cellular level^12^. Similar innovative strategies aimed at bridging the gap between sequence and cell-type specific function are required to tackle these challenges in human genetics.

Cell-types are evolutionary units driven by genomic variation^13^. Similar to speciation, the emergence of new cell-types is a result of changes to gene regulatory networks that drive unique cellular functions or phenotypes in response to evolutionary pressures^14,15^. The stable establishment and maintenance of cell-type specific regulatory programs are broadly regulated by epigenetic mechanisms, which in turn can reflect the evolutionary history of a cell^16,17^. From this perspective, the study of similarities and differences between different cell-types, otherwise known as comparative cell biology^13^, can provide an understanding of the molecular mechanisms underpinning phenotypic diversity at varying evolutionary timescales when comparing information either within or across species. Thus, performing cross-cell-type comparative epigenomic analysis within an individual species could provide information analogous to that obtained from cross-species comparison, offering insight into the evolutionary basis of sequence function and variation at a different scale.

Chromatin is both a product of evolution and a mode of action for evolutionary forces acting on eukaryotic genomes at various scales^18–20^. Polycomb repressive complex 2 (PRC2)-mediated silencing via trimethylation of lysine 27 on histone 3 (H3K27me3) is an ideal example of the co-evolution between chromatin dynamics and genomic variation. While PRC2 is highly conserved across eukaryotes, its H3K27me3-related function has evolved over time in response to evolutionary pressures driving cellular diversification observed in multicellular organisms^21–23^. PRC2 demarcates regions of facultative heterochromatin (fHC) across the genome, which can dynamically adopt open or condensed conformations depending on the spatiotemporal context within a cell. This chromatin plasticity is critical for cells to adapt to different developmental cues, establish distinct gene regulatory programs and maintain specific cellular functions^24^. From an evolutionary perspective, the study of chromatin accessibility and its variation across diverse cell-types within a species could provide insight into the impact of selective constraints on genome function and cellular phenotypic diversity.

Here, we introduce the concept of cellular constraint— wherein the constrained accessibility of a genomic locus across diverse cell-types indicates that the regulatory activity of the locus is functionally required in a cell-type-specific manner. On this basis, cellular constraint measured at a genomic position implies an epigenetically safeguarded locus encoding critical determinants of molecular function. Similar to measures of evolutionary constraint^8^, evaluating patterns of cross-cell-type fHC conservation (rather than cross-species DNA sequence conservation) enables identification of loci with important function in any biological context— agnostic to cell- or tissue-type, developmental stage, trait, or disease. Leveraging this signature of epigenetic conservation, we illustrate the utility of cellular constraint for the scalable and interpretable prioritization of functional loci contributing to cellular phenotypes, traits, and disease across the human genome.

## Results

### Broad domains of H3K27me3 offer a distinctly informative chromatin signature of cell identity regulation

To evaluate patterns of individual histone modifications (HMs) and their regulatory activity across diverse cell- and tissue-types, we analyzed EpiMap reference epigenomes of seven histone modifications (H3K9ac, H3K27ac, H3K4me1, H3K4me3, H3K9me3, H3K27me3, H3K36me3) from 833 biosamples^25^. We compared the breadths of chromatin immunoprecipitation sequencing (ChIP-seq) peaks of different HMs across all 833 biosamples and observed significant variation amongst the top 5% broadest peaks (all pair-wise comparisons; two-tailed Wilcoxon rank sum test, *P* < 2.2 x 10^-16^; **Figure 1A**). We found that genes nearest to the top 5% of broadest HM peaks annotated using ChIPseeker^26^ were largely distinct, with evidence of co-activity between H3K9ac, H3K4me3 and H3K27me3 likely marking bivalent promoters^27,28^ (**Figure 1B**). To delineate these differences, we measured the enrichment of 147 differentiation- and development-related gene ontology (GO) terms in the top 200 genes marked by the broadest HM peaks (**Figure 1C**). We observed that broad H3K27me3 domains were associated with a greater number of significantly enriched GO terms (119 of 147 GO terms tested; *P* ≤ 0.01) and exhibited stronger enrichment compared to other HMs including H3K4me3 and H3K27ac (**Figure 1C)**. This data suggests that broad H3K27me3 domains offer a distinctly informative signature of cellular development and differentiation processes underpinning the genetic regulation of cell identity.

**Figure 1:**
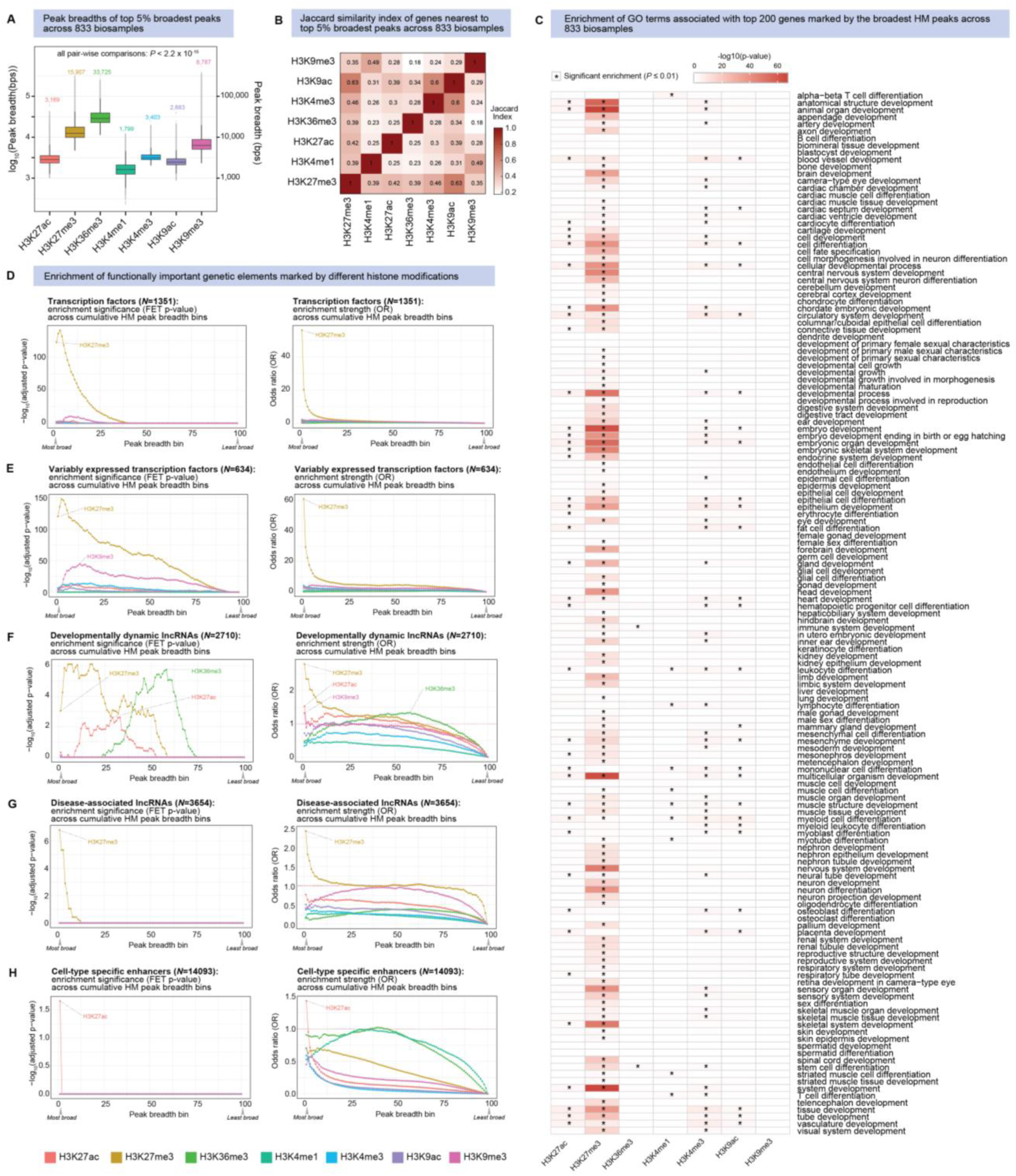
Comparison of histone modification regulatory activity across diverse cell- and tissue-types. **(A)** Comparison of seven different HM peak breadths (bp) of top 5% broadest peaks called across 833 biosamples representing diverse cell- and tissue-types (all pair-wise comparisons; two-tailed Wilcoxon rank sum test, *P* < 2.2 x 10^-16^). Annotated values on boxplot represent average peak breadth for each HM. **(B)** Jaccard similarity index of genes nearest to the top 5% broadest peaks called across 833 biosamples for different HMs. **(C)** Enrichment of cell differentiation and development-related GO terms associated with the top 200 genes marked by the broadest HM peaks across 833 biosamples. **(D-H)** Enrichment of representative sets of functionally important genetic elements marked by different HMs. For each HM, peak breadths overlapping gene-sets are summed across 833 biosamples and cumulatively binned for enrichment analyses. Plots represent significance (Fisher’s exact test; FET P-value; left column) and strength (odd’s ratio; OR; right column) of enrichment of (**D**) transcription factors (TFs) (*N*= 1,351), (**E**) variably expressed TFs (VETFs) (*N*= 634), (**F**) developmentally dynamic lncRNAs (*N*= 2,710), (**G**) disease-associated lncRNAs (*N*= 3,654) and (**H**) cell-type specific enhancers (*N*= 14,093).

We expanded on this to measure the enrichment of other types of functionally important genetic elements marked by HMs patterns across diverse cell-types (**Figures 1D-1H**). To do this, we cumulatively binned loci based on overlapping HM peak breadth summed across 833 biosamples and measured the enrichment of functional gene-sets in each bin. We observed that transcription factors (TFs; *N* = 1,351) were extremely enriched in broad H3K27me3 domains (top bin; *P* = 6.4×10^-142^; odds ratio (OR) = 56.2; **Figure 1D**). Amongst these, a subset of variably expressed TFs (VETFs; *N* = 643) displaying cell-type-specific expression patterns were strongly enriched (top bin; *P* = 3.5×10^-148^; OR = 61.3; **Figure 1E**), which aligned well with our previous findings^29^. Broad domains of H3K9me3 were also enriched for TFs (top bin; *P* = 5.2×10^-11^; OR = 2.3) and VETFs (top bin; *P* = 1.5×10^-45^; OR = 3.9), though this enrichment was substantially weaker compared to broad H3K27me3 (**Figure 1E**).

Next, we evaluated non-coding regulatory elements and found that developmentally dynamic long non-coding RNAs^30^ (lncRNAs) (*N* = 2,710) were enriched in broad domains of H3K27me3 (top bin; *P* = 7.8×10^-7^; OR = 2.8) and H3K27ac (top bin; *P* = 0.002; OR = 1.5), as well as moderate domains of H3K36me3 (top bin; *P* = 1.8×10^-6^; OR = 1.3) (**Figure 1F)**. However, only broad H3K27me3 showed significant enrichment of disease-associated lncRNAs (*N* = 3,654) (top bin; *P* = 1.5×10^-7^; OR = 2.4; **Figure 1G**). We also observed that only broad H3K27ac showed enrichment of cell-type-specific enhancers (*N* = 14,093) displaying enhancer activity in only 1 out of 295 cell-types in the EnhancerAtlas database^31^, which is concordant with previously reported links to super-enhancer activity^32–35^ (top bin; *P* = 0.02; OR = 1.4; **Figure 1H**). Overall, these data suggest that broad H3K27me3 domains are uniquely enriched for both functionally important coding and non-coding elements with cell-type specific regulatory activity and this association weakens as H3K27me3 peak breadth narrows.

### Cross-cell-type facultative heterochromatin conservation reveals genomic regions of cellular constraint

Given that broad H3K27me3 is a hallmark of fHC formation known to selectively repress important genes required upon spatiotemporal developmental cues^24,28,36,37^ (**Figures 1C-1G**), we next sought to characterize fHC regulatory activity across the genome by performing cross-cell-type comparative epigenomic analysis. Evaluating H3K27me3 ChIP-seq peak alignment across 833 human biosamples from EpiMap^25^, we observed that patterns of broad fHC were conserved at specific sequences across diverse cell- and tissue-types (**Figure 2A).** This chromatin signature of selectively conserved fHC repression was observed at a subset of loci known to be associated with cell-type specific regulation such as *MYOD1* and *H19*, and not found at housekeeping and structural genes such as *GAPDH* and *MYH2* for example (**Figure 2B)**. Further, this pattern of deposition was not observed for other histone marks, including H3K9me3, another repressive HM (**Figure S1A**).

**Figure 2:**
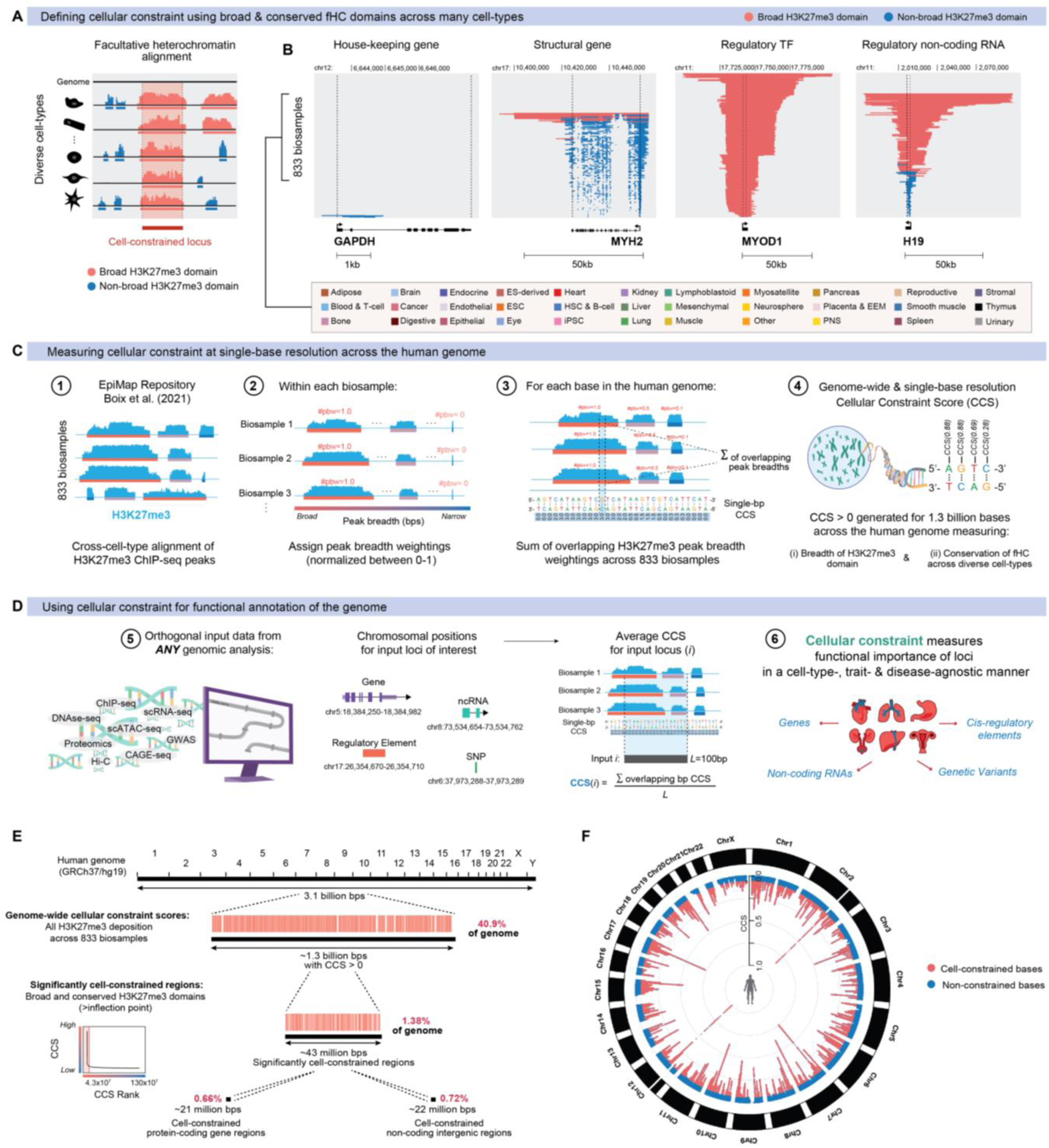
Overview of cellular constraint across the human genome. (**A-B**) Defining cellular constraint. **(A)** Evaluating facultative heterochromatin (fCH) alignment across diverse cell-types identifies genomic loci with restricted accessibility, whose activity is functionally required in specific spatiotemporal contexts. **(B)** Examples show fCH domains, marked by broad H3K27me3 conserved across many cell-types, selectively repress loci with cell-type specific regulatory function. Plots display aligned ChIP-seq peaks from 833 EpiMap biosamples, showing H3K27me3 deposition over a locus stacked vertically and sorted on breadth. Domains ranked in the top 5% within each biosample are considered broad (red) and the remainder non-broad (blue). (**C-D**) Measuring and using cellular constraint. **(C)** Schematic of single-base cellular constraint score (CCS) calculation: For each base-pair in the human genome, the (i) breadth and (ii) conservation of H3K27me3 deposition is measured across diverse cell- and tissue-types. A genome-wide CCS metric is derived by summing overlapping H3K27me3 domain breadth weightings from 833 EpiMap biosamples at single-base resolution. **(D)** Orthogonal input data from any genomic analysis can be interfaced with the CCS as a weighting metric to evaluate functional loci. (**E-F**) Genome-wide coverage of cellular constraint. (**E**) The CCS reference metric covers 41.4% (1.3 billion bases; CCS > 0) of the human genome, of which a subset of 1.4% (43 million bases) are defined as significantly cell-constrained domains. **(F)** Circular Manhattan plot of genome-wide distribution of CCS values.

Sequences marked by cross-cell-type conserved fHC display restricted accessibility within the limited cell-type(s) and/or developmental context(s) in which their expression/activity is spatiotemporally required. Conversely, these loci are broadly inaccessible in cell-types and contexts in which their ectopic activity is actively silenced via fHC formation to maintain transcriptional stability within a given lineage (**Figures S1B and S1C**). We define these sequences, likely to have cell-type specific regulatory function based on epigenetically constrained chromatin accessibility across different cell- and tissue-types, as regions under cellular constraint.

### Defining cellular constraint at single-base resolution

Next, we set out to quantify cellular constraint at single-base resolution by measuring the breadth and conservation of fHC deposition across diverse cell- and tissue-types across the human genome. Using reference epigenomes from 833 EpiMap biosamples^25^, covering 33 major tissue groups and different life stages (embryonic, newborn, child, adult), we first performed cross-cell-type H3K27me3 peak alignment. Within each biosample, we ranked H3K27me3 peaks by breadth and normalized these values between 0-1 to generate peak breadth weightings, where a value of 1 represents the single broadest peak in a given biosample. For each base in the genome, we then summed overlapping peak breadth weightings across all 833 biosamples and normalized these values between 0-1 to generate genome-wide cellular constraint scores (CCS). These CCS values represent a metric of fHC domain breadth and conservation across diverse cell- and tissue-types measured at each genomic position across the human genome (**Figure 2C and see Methods**). As cellular constraint measures functional importance in a cell-type, trait, or disease-agnostic manner, it can theoretically be applied to evaluate *any* locus of interest in *any* biological context (**Figure 1D**).

We calculated CCSs > 0 for 1.3 billion bases (41.4%) in the human genome (hg19; GRCh37 and hg38; GRCh38 available, **see Methods**). The remaining 58.6% of the genome displayed no detectable H3K27me3 signal in any biosample and were assigned a value of 0. We compared two methods to determine a significance threshold by using an empirically determined statistical cut-off (CCS ≥ 0.03; 81,023,692 bases) or using the inflection point of the CCS curve (CCS ≥ 0.06; 43,240,489 bases) (**Figure S2A**). We decided to use the more stringent inflection point cut-off to call significantly cell-constrained bases, as the CCS distribution was right-skewed and the enrichment of functional gene sets was comparable between methods (**Figure S2B**). The average breadth of significantly cell-constrained domains (blocks of consecutive bases with the same CCS value) was ∼60bp in length ranging from min 24bp to max 42,600bp.

To test whether cell- or tissue-type sampling bias within the 833 EpiMap biosamples influences CCS calculation, we performed three complementary analyses. First, we performed bootstrapping analysis via random resampling of biosamples (*N*= 10 permutations) (**Figure S2C**). Second, we iteratively removed all biosamples from each one of the 33 tissue-groups represented (*N*= 32 permutations) (**Figure S2D).** We found that both permutation approaches did not significantly alter CCS calculations or inflection point threshold. Last, we compared CCS values generated from EpiMap^25^ (833 biosamples) versus Roadmap^38^ (111 biosamples) and found that the CCS remained stable across the genome (Spearman’s ρ = 0.92; *P*<2.2×10^-16^) (**Figure S2E**). Together these data suggest there is sufficient cellular diversity sampled in publicly available epigenomic databases to stably estimate cellular constraint in humans.

Using the inflection point of the CCS curve, we define 43 million bases or 1.4% of the human genome as significantly cell-constrained (**Figures 2E and 2F**). Briefly, we found that cell-constrained bases were distributed equally across both genic (7.7% exonic and 42.6% intronic) and intergenic (44.8% intergenic, 2.9% 3’ UTR and 1.8% 5’ UTR) regions (**Figure S2F**). The vast majority of cell-constrained bases corresponded to non-coding sequences spanning intronic regions within gene bodies, *cis*-regulatory elements nearby gene promoters as well as distal regulatory elements located more than tens of kilobases away from their nearest protein-coding gene (**Figures S2G-S2H**).

### Conservation of cross-cell-type H3K27me3 patterns across distant species

The conservation of PRC2 function in eukaryotes points to the critical function of H3K27me3 in preserving domains of heterochromatin, specifically fHC, through evolution^23,39^. We therefore tested whether domains of cross-cell-type H3K27me3 were also conserved across species. To do this, we calculated genome-wide CCS values for the mouse genome (mm10; GRCm38) using H3K27me3 ChIP-seq data from 181 ENCODE mouse biosamples^40^ (**Figure S3A**). Comparing CCS values for overlapping syntenic blocks of 100bp between human and mouse genomes (*N* = 2,692,660), we found that CCS metrics calculated between species were positively correlated (**Figure 3A**; Spearman’s ρ=0.59; *P*<2.2×10^-16^). This suggests that cell-type-specific and/or context-dependent patterns of H3K27me3 at both coding and non-coding sequences are epigenetically conserved between these species, separated by ∼80 million years of evolution. These results are consistent with previous findings of cross-species histone mark conservation^39,41^, indicating that variation in H3K27me3 might be an evolutionary mechanism underlying regulatory divergence between species. While we found species-specific H3K27me3 patterns for a subset of homologous genes (**Table S2**), the CCS values of human-mouse homologs were largely positively correlated (**Figure S3B**; Spearman’s ρ=0.68; *P*<2.2×10^-16^). Together, these results suggest that cross-cell-type repressive fHC domains marking cell-constrained sequences are largely conserved and maintained in distantly related mammalian species.

**Figure 3:**
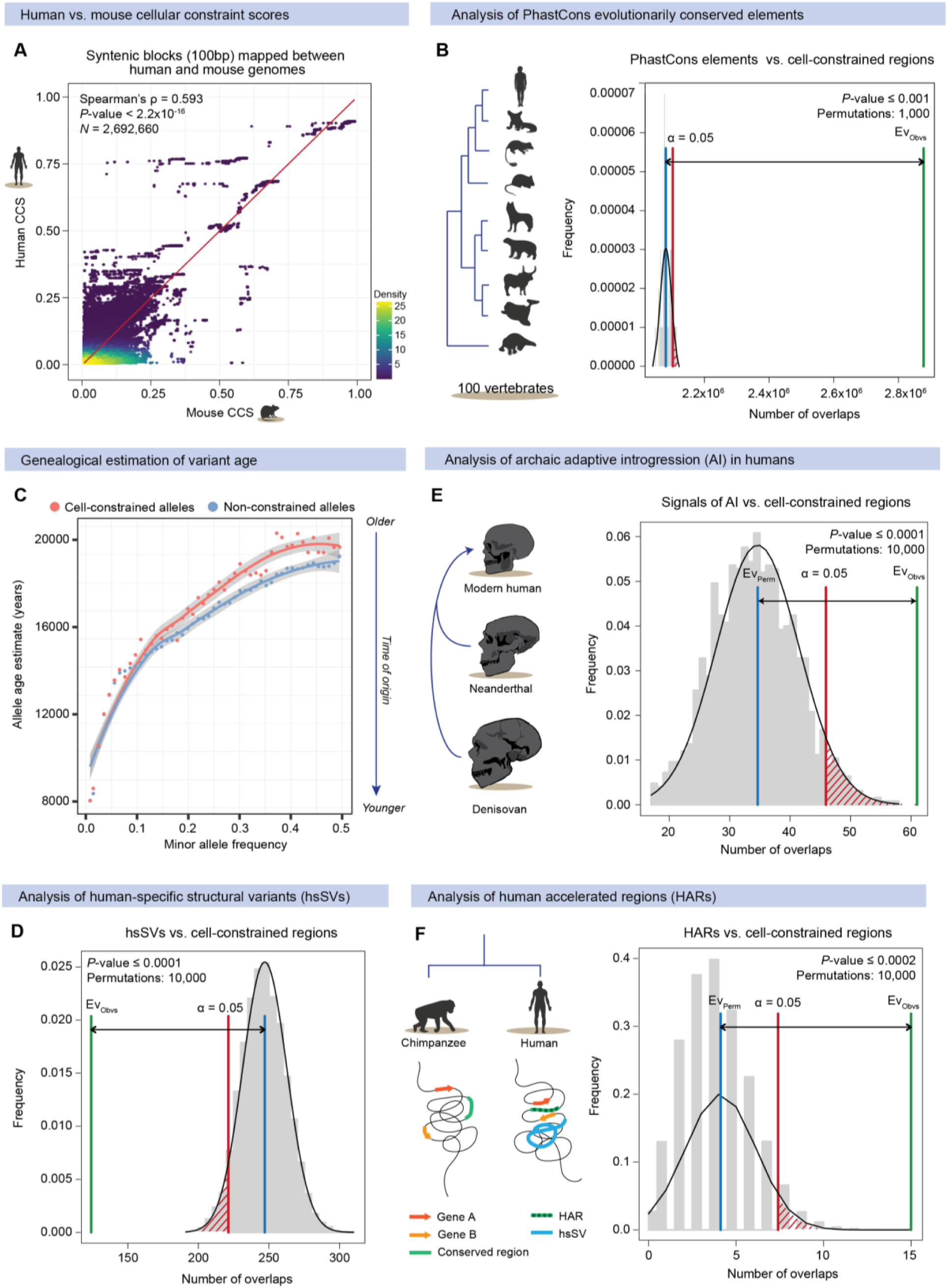
Enrichment of signals of evolutionary conservation and adaptation at cell-constrained bases. **(A)** Correlation between human and mouse cellular constraint calculated for 100bp syntenic blocks mapped between the human genome (hg19) and the mouse genome (mm10) (Spearman’s ρ = 0.593; *P*<2.2×10^-16^, Wilcoxon’s rank sum test). **(B)** Permutation test evaluating overlap between cell-constrained bases and PhastCons conserved elements (*P* ≤ 0.001) based on *n* = 1,000 random permutations. **(C)** Allele age estimate relative to allele frequency of cell-constrained variants and non-constrained variants. GEVA, genealogical estimation of variant age. **(D-F)** Permutation tests evaluating overlap between cell-constrained bases and **(D)** human-specific structural variants (hsSVs) (*P* ≤ 0.0001), **(E)** Signals of archaic adaptive introgression (*P* ≤ 0.0001) and **(F)** human accelerated regions (HARs) (*P* ≤ 0.0002) based on *n* = 10,000 random permutations. The average expected number of overlaps (blue) is compared against the observed number of overlaps (green), with α = 0.05 confidence interval indicated (red).

### Cellular constraint enriches for conserved loci conferring evolutionary advantage

Next, we sought to directly evaluate the association between measures of cross-cell-type fHC conservation (cellular constraint) and cross-species DNA sequence conservation (evolutionary constraint). We first tested whether cell-constrained domains were enriched for PhastCons^42^ elements across 100 vertebrate species (PhastCons score cut-off ≥ 0.9; 12.4×10^7^ nucleotides). Using genome-wide permutation analysis^43^, we found significant co-localization of cell-constrained bases and PhastCons evolutionarily conserved bases (**Figure 3B**; *P* ≤ 0.001; 1,000 permutations). This enrichment was independently observed in cell-constrained non-coding regions (**Figure S3C;** *P* ≤ 0.001; 1,000 permutations), confirming this signal is not inflated due to the proximity of H3K27me3 domains to coding sequences. While we observe significant co-localization of cell-constrained and evolutionarily-constrained bases, we found that these metrics differentially prioritize sets of genetics elements (**Figure S4A and S4B**). Comparing enrichment of GO terms associated with the top 100 genes prioritized by cellular constraint (CCS) versus evolutionary constraint (PhastCons mammal 470-way), we found that cell-constrained genes enriched for biological processes involved in cell fate commitment and organ system development whereas evolutionarily-constrained genes enriched for metabolic processes (**Figure S4C**). Direct comparison of the enrichment of gene-sets involved in cell-type specific regulation (VETFs and TFs), cell fate specification and organ morphogenesis showed that cellular constraint significantly enriched for these subsets to a greater extent than evolutionary constraint (**Figure S4D-S4F**).

Moreover, we found that the estimated allele age^44^ of cell-constrained common variants was significantly older compared to non-constrained common variants (allele frequency range ∼0.25-0.75), indicating that the time of origin of mutations falling in cell-constrained domains date further back in evolutionary history (**Figure 3C)**. The distribution of cell-constrained and non-constrained alleles is comparable across different minor allele frequency (MAF) bins (0-0.5), indicating no significant bias in the prevalence of alleles in these regions (**Figure S3D**). We also observed significant under-enrichment of human-specific structural variants (hsSVs) overlapping with cell-constrained domains (**Figure 3D**; *P* ≤ 0.0001; 10,000 permutations), suggesting these regions are intolerant to change. Taken together, these data strongly suggest that purifying selection is acting on regions of cellular constraint to preserve their ancestral, functional state over evolutionary time.

Next, we sought to assess whether cell-constrained loci have also undergone positive selection over time to drive adaptive evolution in human populations. Modern humans interbred with archaic hominins including Neanderthals and Denisovans resulting in present-day human genomes containing introgressed genomic regions, which conferred surviving populations with an evolutionary advantage via a process known as adaptive introgression^45–48^. We observed significant enrichment of signals of archaic adaptive introgression in the human lineage within cell-constrained domains based on permutation sampling (**Figure 3E**; *P* ≤ 0.0001; 10,000 permutations).

Human accelerated regions (HARs) are sequences that, despite being evolutionary conserved in other species, have acquired high numbers of nucleotide substitutions in human genomes since divergence from our common ancestor with chimpanzees^49,50^. Surprisingly, we also found significant co-localization of cell-constrained loci and HARs (**Figure 3F**; *P* ≤ 0.0002; 10,000 permutations). Together, these analyses show that the 1.4% of the genome safeguarded by cellular constraint are enriched for loci that have been conserved across millions of years of evolution and been retained in modern humans for selective advantage. This suggests that these regions contain loci governing critical functions underpinning cell and organismal phenotypes.

### Prioritization of functional lncRNAs governing development and disease

Evolutionary conservation is strongly linked to the functionality of coding and non-coding sequences^51,52^. Given that signals of evolutionary constraint are enriched in cell-constrained regions, we evaluated its association with lncRNA function. We first measured enrichment of cell-constrained lncRNAs among lncRNAs conserved across seven vertebrate species representing various evolutionary distances^30^ (rhesus macaque: 25 million years ago (Mya); mouse: 90 Mya; opossum 180 Mya; chicken: 300 Mya). We observed increasing enrichment of cell-constrained lncRNAs represented in older evolutionary age groups (**Figure S5A**; *P<*0.01, two-tailed Fisher’s exact test). Enrichment of cellular constraint was also seen across twelve different primate species^53^ (**Figure S5B;** *P<*0.01) and among developmentally dynamic lncRNAs^30^ variably expressed through early organogenesis to adulthood in seven major organ systems (**Figure S5C;** *P=*4.63×10^-29^).

We generated CCS values for 127,802 transcripts from LNCipedia^54^ and used the inflection point of the CCS curve to define a set of 5,723 significantly cell-constrained lncRNAs (**Figure S5D** and **Table S3**). Cellular constraint prioritized many functionally validated developmental and disease regulators including *HOTAIR*^55^*, FENDRR*^56^, *H19*^57^*, lnc-WISP2-2*^58^ and *PANCR*^59,60^ (**Figure S5E).** Moreover, we observed significant enrichment of lncRNA-disease associations across three independent databases including LncTaRD^61^ (*P=*4.71×10^-7^, two-tailed Fisher’s exact test), LncRNADisease^62^ (*P=*4.85×10^-9^) and MNDR^63^ (*P=*5.13×10^-7^) (**Figure S5F**). Collectively, these results suggest that cellular constraint is a strong indicator of lncRNA functionality.

### Cell-type specific prediction of non-coding loci regulating cellular phenotypes

When integrated with complementary data indicating a locus’ activity (e.g. transcript abundance) or association (e.g. genome-wide significance level) in a given developmental or disease context, cellular constraint can directly inform cell-type-specific function or molecular mechanism (**see Methods**). As proof-of-concept validation, we analyzed bulk cap analysis of gene expression sequencing (CAGE-seq) of a time-course of *in vitro* human induced pluripotent stem cell (hiPSC) cardiac differentiation (**Figure S6** and **Supplementary Information**).

Using cellular constraint to weight transcript abundance at each differentiation timepoint (**Figure S6** and **Table S4**), we identified *NKX2-6-AS1* (*RP11-175E9.1*; ENST00000523874.1) as a CCS-prioritized candidate novel divergent lncRNA transcript predicted to regulate cardiogenesis (**Figure S6B-S6G**). We functionally perturbed *NKX2-6-AS1* transcription in hiPSCs via targeted CRISPRi and found that its loss-of-function resulted in failure to differentiate cardiac troponin T^+^ cardiomyocytes *in vitro* (**Figure S6H and S6I**). Transcriptional profiling data suggest that *NKX2-6-AS1* maintains cardiac fate upon divergent initiation of *NKX2-6* transcription during early stages of cardiac specification (**Figure S6J-S6O)**. This illustrates that cellular constraint scores provide an effective weighting metric to assist with the interpretation of context-specific locus activity and function in regulating cellular phenotypes.

### Prioritization of genetic variants contributing to complex traits and disease across the human genome

Measures of evolutionary constraint and biological function have been used to identify genetic variants underlying disease susceptibility and complex trait phenotypes^5,7,64^. Since cell-constrained domains are enriched for loci with high levels of sequence conservation and molecular function, we hypothesized that single-base CCS values could be used to systematically prioritize genetic variants associated with complex traits and disease across the human genome.

Most trait- and disease-associated variants are non-coding, and are thus most likely to influence disease risk by altering gene expression or splicing^65,66^. As such, we analyzed GTEx expression quantitative trait loci (eQTLs) which represent variants influencing gene expression levels in different tissues from 449 individuals^67^. We generated CCS values for 2.1 million GTEx *cis*-eQTLs across 49 cell- and tissue-types^67^ and used the inflection point of the CCS curve to define a set of 99,010 significantly cell-constrained eQTLs (**Figure 4A** and **Table S5**). Across all cell- and tissue-types, we observed that cell-constrained eQTLs had significantly lower p-values, larger effect sizes, and lower minor allele frequencies (each with *P* < 1×10^-300^, Wilcoxon’s Rank Sum Test), compared to non-constrained eQTLs (**Figure 4B and 4C)**. The observation of larger effect sizes at rarer variants is consistent with a model of negative selection, suggesting that cell-constrained eQTLs are more likely to be detrimental in humans and therefore suppressed at low frequencies in the population.

**Figure 4:**
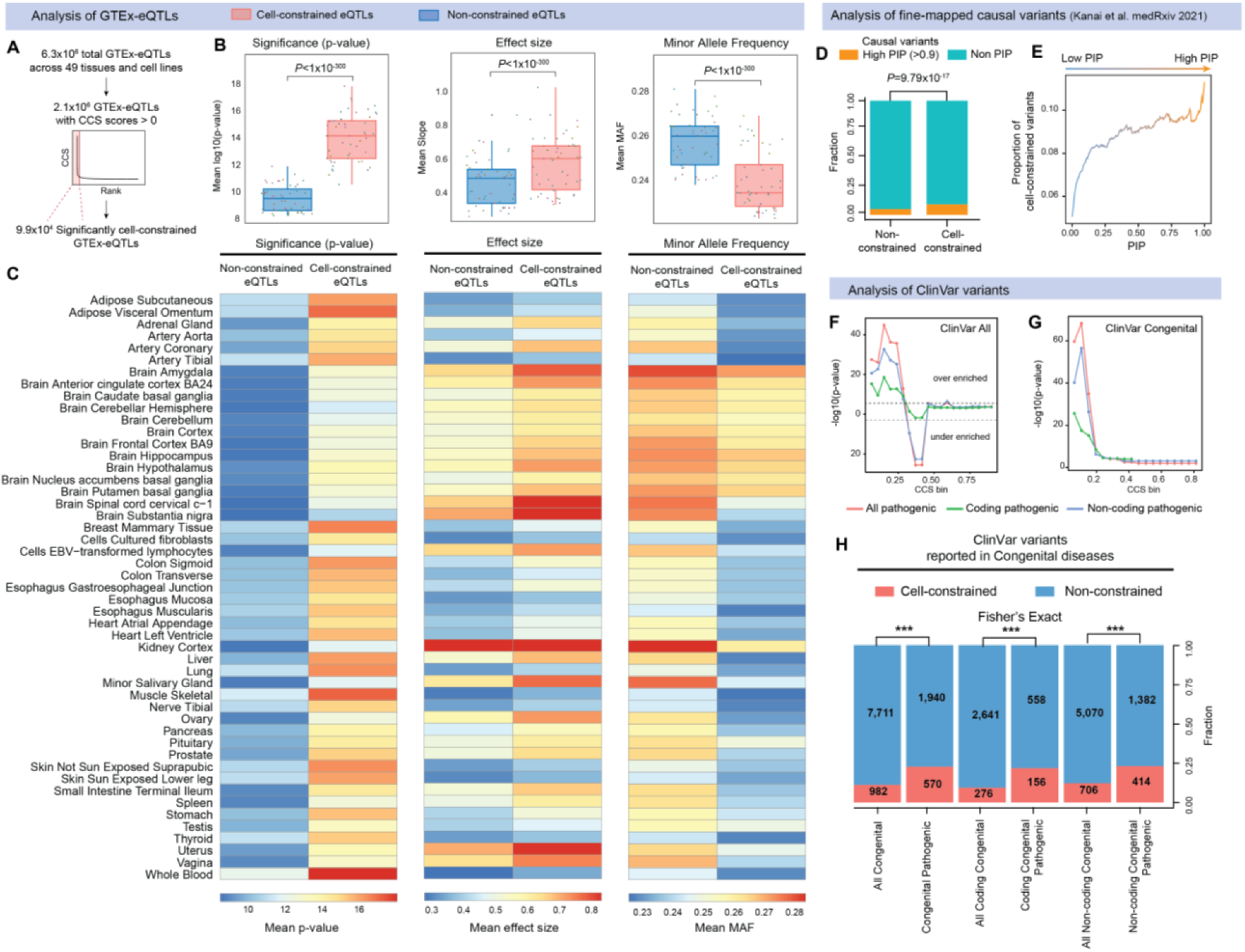
Cellular constraint prioritizes genetic variants linked to complex traits and disease. **(A-C)** Cellular constraint-based prioritization of GTEx-eQTLs across 49 tissues and cell lines. (**A**) CCS generation for all GTEx-eQTLs to define a set of significantly cell-constrained eQTLs (*N* = 99,010). (**B-C**) Mean significance, slope, and minor allele frequency of cell-constrained eQTLs (*N* = 99,010) versus non-constrained eQTLs (*N* = 2,198,997) across 49 GTEx tissues (*P*<0.001, two-sided Wilcoxon’s signed-rank test). **(D-E)** Cellular constraint-based prioritization of fine-mapped variant trait-pairs from Kanai *et al.* **(D)** Fraction of cell-constrained variants in high PIP (>0.9) and non-PIP variant categories (*P*=9.79×10^-17^, two-tailed Fisher’s exact test) **(E)** Proportion of cell-constrained variants enriched across PIP variant bins. (**F-H)** Cellular constraint-based analysis of ClinVar variants. (**F**) Significantly cell-constrained variants are enriched for pathogenic ClinVar variants across all diseases reported and (**G**) display stronger enrichment for pathogenic variants linked to congenital disorders (one-tailed Fisher’s exact test). This enrichment attenuates in extremely high CCS regions (CCS ∼>0.2), suggesting that pathogenic variants are less tolerated as CCS increases. (**H**) Fraction of cell-constrained variants for Clinvar variants associated with congenital diseases in subsets of pathogenic, coding pathogenic and non-coding pathogenic variant categories (\*\*\**P*<0.001, two-tailed Fisher’s exact test).

Fine-mapping methods help identify causal variants for a trait or disease from genome-wide association study (GWAS) hits^68^. GWAS identified loci are often challenging to interpret due to confounding issues such as strong linkage disequilibrium (LD) at a given locus. We sought to evaluate whether variants independently prioritized by cellular constraint were enriched for fine-mapped variants underlying a range of complex traits. For this, we assessed an atlas of 4,518 high confidence (posterior inclusion probability or PIP > 0.9) putative causal variants ^69^ using fine-mapping across 148 complex traits in three large-scale biobanks (BioBank Japan, FinnGen and UK BioBank). We found that cell-constrained variants were significantly enriched for putative causal variants compared to non-constrained variants (*P*=9.79×10^-17^; two-tailed Fisher’s exact test) (**Figure 4D**). Further, we observed a significant increase in the proportion of cell-constrained variants represented in increasingly higher PIP variant groups, indicating that mutations in high CCS loci are more likely to be of clinical significance (**Figure 4E**). These data show that genetic variants falling in cell-constrained domains are more likely to have causative effects on complex traits and diseases, suggesting that the CCS metric can predict the functional relevance of both coding and non-coding variants.

Lastly, we tested whether variants prioritized by cellular constraint were enriched for rare clinically relevant variants with evidence of pathogenicity in ClinVar^70^. We generated CCS values for 501,715 ClinVar variants and used the inflection point of the CCS curve to define a set of 40,650 significantly cell-constrained variants. We found that high CCS variants were significantly enriched for pathogenic ClinVar variants across all diseases reported (*P*=1.12×10^-26^, two-tailed Fisher’s exact test) and this enrichment was seen for both coding and non-coding variants (*P*=1.09×10^-13^ and *P*=1.92×10^-19^, respectively) (**Figure 4F**). Notably, focusing on congenital diseases we found that cell-constrained variants showed even stronger enrichment for pathogenic ClinVar variants reported in this subset of conditions affecting early fetal development (all *P*=1.18×10^-59^, coding *P*=2.99×10^-23^ and non-coding *P*=8.59×10^-39^; two-tailed Fisher’s exact test) (**Figure 4G and 4H**). Mutations affecting developmental gene programs, especially those in core transcription factors, are major contributors to congenital disorders^71–73^ hence this enrichment is expected given that cell-constrained domains are strongly associated with regulators of cell identity. Together, these data suggest that variants prioritized by cellular constraint are more likely to be pathogenic and be of clinical significance.

### Cell-constrained variants have stronger influence on the SNP-based heritability and polygenic prediction of complex traits and diseases

To assess the impact of variation within cell-constrained domains on the genetic architecture of human diseases and complex traits, we performed. We used GWAS summary statistics across 28 complex traits and diseases from UK BioBank^74^. We found that across the traits tested, cell-constrained SNPs had on average significantly higher per-SNP heritability enrichment compared with non-constrained SNPs (**Figure 5A**). Cell-constrained SNPs were enriched in both the proportion of causal variants and the magnitude of effect sizes (**Figure 5B**). Next, we compared the effect of different functional annotations on polygenic risk score (PRS) prediction for the same 28 UK BioBank^74^ complex traits and diseases. We found that using cellular constraint as a continuous (CCS annotation for all SNPs) or binary (CCS annotation for only significantly cell-constrained SNPs) model improved the prediction accuracy compared to the no annotation model (**Figure 5C**). Moreover, we found that the combined annotation using CADD^75^, 29-way mammalian constraint^76^, and binary CCS performed the best of all the models tested, indicating that cellular constraint is complementary to existing genomic annotations and captures unique biological information not accounted for by other annotations (**Figure 5C and 5D**). Together, these data demonstrate that causal variants with large effect sizes on complex traits and diseases are more likely to fall within domains of cellular constraint, suggesting that high-CCS loci underlie major phenotypic determinants.

**Figure 5:**
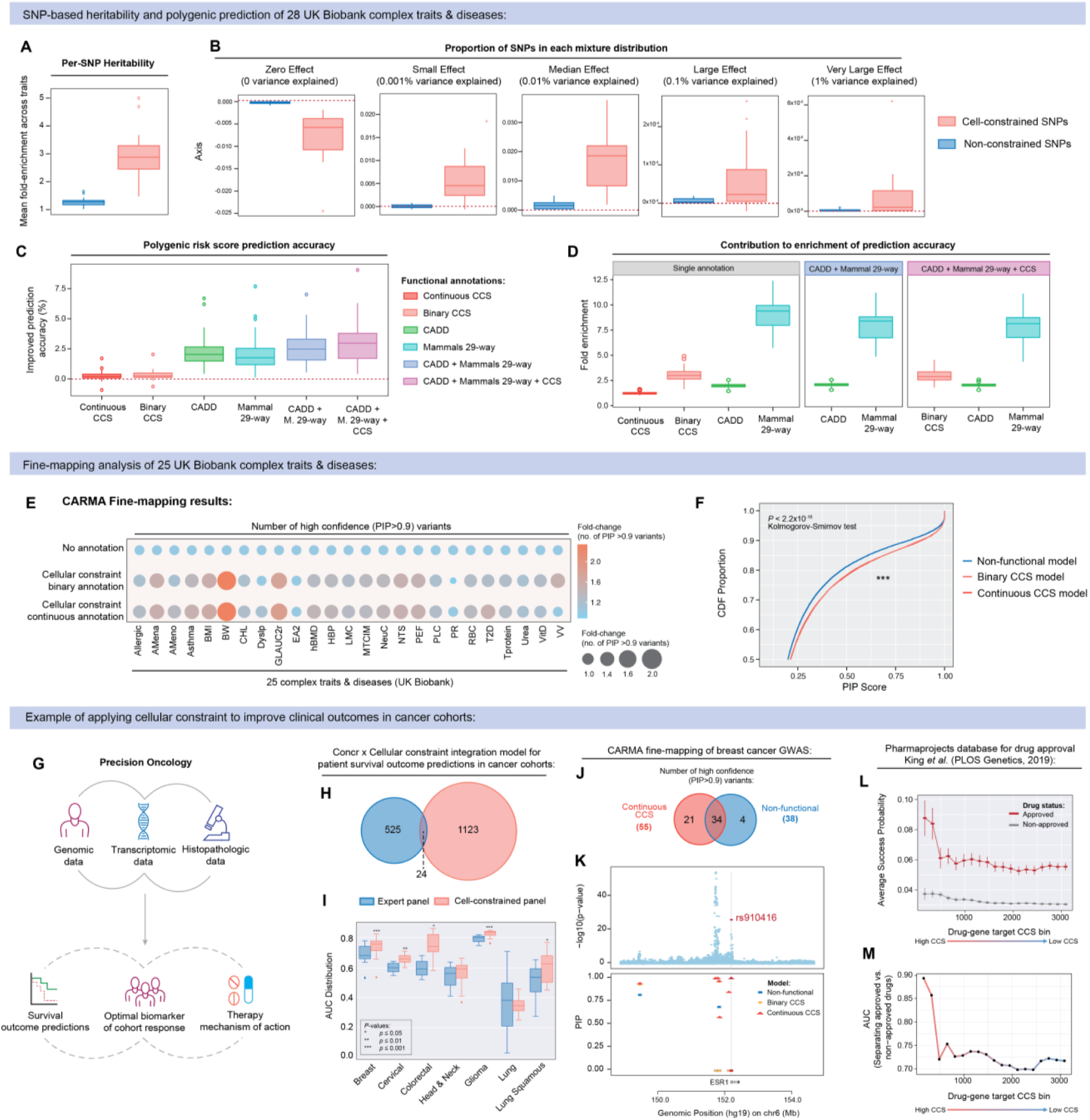
Leveraging cellular constraint to improve clinically translatable outcomes. (**A-D**) Annotation stratified SBayesRC analysis comparing cell-constrained SNPs vs. non-constrained SNPs using GWAS summary statistics of 28 complex traits and diseases from UKBiobank ^74^. Meta-analysis results across traits for parameter enrichment measuring (**A**) Per-SNP Heritability and **(B)** Effect size in various groups. **(C)** Comparison of different genomic annotation models and their effect on percentage improvement in polygenic risk score prediction accuracy. **(D)** Contribution to the enrichment of prediction accuracy for each annotation model. Continuous CCS = all SNPs; Binary CCS = only significantly cell-constrained SNPs; CADD = Combined Annotation-Dependent Depletion SNPs ^75^; M. 29-way = 29-way mammalian constrained SNPs ^76^. (**E-F**) Functionally informed fine-mapping of 25 UKBiobank complex traits and diseases using cellular constraint as a binary or continuous model of functional annotation. **(E)** Number of high confidence (PIP >0.9) fine-mapped variants identified for each trait and disease. **(F)** Cumulative distribution function (CDF) of PIP scores using different models of functionally informed fine-mapping. One-way Kolmogorov-Smirnov tests show that CDFs for PIP scores obtained from both cellular constraint annotation models (light and dark red) are higher than those of the non-functional baseline model (blue) (\*\*\**P*< 2.2×10^-16^). (**G**) Schematic of clinical therapeutic pipelines for early diagnosis, prognosis, and drug discovery in precision oncology. **(H)** Variant panels selected via expert curation (*N=*525 variants) vs. cellular constraint (*N=*1,123 variants). **(I)** Prediction accuracy of prognostic machine learning model for the prediction of 5-year survival outcomes in different cancer cohorts using cell-constraint vs. expert curation annotation. (**J)** Fine-mapping of breast cancer GWAS meta-analysis ^79^ using cellular constraint as a continuous model of functional annotation. **(K)** Example of cell-constrained fine-mapped variant rs910416. Plot displays GWAS *P*-values (top) and corresponding PIP scores under different annotated fine-mapping models represented by unique shapes (bottom). **(L)** Average success probability of drugs-gene-indication trios reported in King *et al.* (2019)^82^ from the Informa pharmaprojects database stratified by gene CCS and approval status. (**M**) Performance accuracy of classifying drug approval for clinical use. Trios are ranked and binned from high to low (left to right) based on the cellular constraint scores of a drug’s target gene.

### Cellular constraint scores improve fine-mapping analysis

We tested whether leveraging cellular constraint as a genomic annotation could improve the power of functionally informed fine-mapping of causal variants associated with complex traits. We compared fine-mapping results from CARMA^77^ without annotations (nonfunctional model) against that with cellular constraint annotation using the CCS as a continuous or a binary variable on 25 well-powered UK Biobank complex traits and diseases. We observed a significantly higher number of high confidence putative causal variants (PIP > 0.9) at cell-constrained sites across traits when using CARMA with either the continuous or binary CCS annotation model (1,553 and 1,487 putative causal variants, respectively) compared to the non-functional model (1,139 putative causal variants) (**Figure 5E and 5F**). Overall, these results indicate that functional annotation of the genome with cellular constraint can improve the power of identifying causal variants underlying complex traits and diseases.

### Using cellular constraint in machine learning for clinical prediction

We next tested the utility of cellular constraint metrics integrated into clinical data analysis pipelines. We focused on models assisting in early patient diagnosis, prognosis and drug discovery that are driven by increasing availability of patient- or disease-specific multi-omic data (**Figure 5G**). First, we evaluated whether integrating cellular constraint annotation into a prognostic machine learning model^78^ could enhance the prediction of 5-year survival outcomes in cancer cohorts. We used a hierarchical Bayesian inference framework to combine domain-specific machine learning sub-models processing multi-omic oncogenic data from patients (DNA sequencing and histopathology) and *in vitro* models (single cell-RNA sequencing)^78^. To test the efficacy of using cellular constraint compared to the clinical gold standard annotation, we evaluated the performance of the survival model comparing an expert curated variant panel (525 variants) to an unsupervised cell-constrained variant panel (1,123 variants), generated by selecting variants above the inflection point of the CCS-weighted curve (**Figure 5H**). These variant panels showed minimal overlap. We found that cellular constraint variant annotation significantly improved the accuracy of 5-year survival outcome predictions across multiple cancer subtypes compared to the clinical gold standard annotation (**Figure 5I)**.

### Fine-mapping example in breast cancer GWAS loci

Next, we illustrate the utility of cellular constraint annotation in identifying putative causal variants underlying diseases that might be overlooked in the absence of this additional functional information. We performed CARMA fine-mapping analysis on 32 breast cancer risk loci identified from a large GWAS meta-analysis of 113,384 breast cancer patients and 113,789 controls of European ancestry^79^. The result demonstrated a significantly higher number of high confidence variants (PIP > 0.9) when using the continuous and binary CCS annotation models (55 and 58 causal variants, respectively), compared to the nonfunctional model (38 causal variants) (**Figure 5J**). Notably, we found that each CCS annotation model identified distinct sets of causal variants which were not shared between them (**Figure 5J**).

We highlight rs910416, which falls in a cell-constrained domain with substantial H3K27me3 repression across many cell-types, leading to a higher PIP when using the continuous CCS model (PIP= 1) compared to the nonfunctional model (PIP = 0) (**Figure 5K**). rs910416 is an intronic non-coding variant in the Estrogen Receptor 1 (*ESR1*) locus and has a causal and experimentally validated association with breast cancer^80^. Interestingly, it has been shown that rs910416 is located in a putative silencer element and that the allele-specific binding of *MYC* introduced by this SNP disrupts the silencer’s function in regulating *ESR1* and *RMND1* expression, leading to breast cancer development^80^. This example illustrates how cellular constraint improves fine-mapping and can be leveraged to discover regulatory mechanisms linking variation to function.

### Leveraging cellular constraint as a predictor of drug approval

While drug approval rates are notoriously low, it has been shown that human genetic evidence linking a drug’s target gene to the condition it treats can double the likelihood of approval for therapeutic use^81^. Given that cellular constraint enriches for causal genetic variants underlying complex traits and disease, we investigated whether the CCS of a drug’s gene target could be used as a predictor of drug approval. To test this, we utilised the Informa Pharmaprojects dataset containing indication-drug-gene trios, their approval status for clinical use and predicted success probabilities from King *et al.* (2019)^82^. We found that cellular constraint can effectively augment the predicted success probability of drugs, as shown by greater separation in average success probability between approved and non-approved drugs with high CCS gene targets (**Figure 5L**). This enables the prediction of clinical success probability (approval status) with high accuracy (AUC) for drugs with high CCS gene targets (**Figure 5M**). These data suggest that cellular constraint annotation of indication-drug-gene trios improves the accuracy of drug success probability predictions and the CCS can act as a predictor of drug approval to inform the development of new therapies.

## Discussion

This study defines cellular constraint as a new genomic annotation. Cellular constraint captures a highly concentrated view of the regulatory genomic landscape, by focusing on 1.4% of the human genome safeguarded by broad domains of fHC conserved across diverse cell-types. We found this repressive chromatin signature to be largely conserved in the mouse genome indicating preservation across an evolutionary span of at least ∼80 million years. While negative selection pressure counteracting variation in cell-constrained regions is apparent, they also appear to be a battleground for active evolutionary forces driving adaptation in recent human evolution. The paradoxical signals of purifying selection and human acceleration suggests that the selective pressures acting on cell-constrained sequences can switch from negative to positive depending on the evolutionary advantage conferred within a population^83^. Further investigation into the extent to which cross-cell-type H3K27me3 patterns are conserved or divergent across species of varying phylogenetic distances could offer valuable insight into epigenetic mechanisms driving evolutionary novelty. Single-base resolution cellular constraint scores allow efficient identification of functionally important coding and non-coding elements across the genome, enabling insight into the impact of genetic variation on cellular phenotypes, human diseases, and complex traits. We highlight the utility of cellular constraint as a functional annotation to enhance the identification of candidate causal variants in both rare and common diseases, improving clinical diagnostic predictions, and prioritizing candidates in drug discovery pipelines.

## Methods

### Evaluating genomic features marked by different histone modifications

For this analysis, we used observed (excluding imputed) chromatin immunoprecipitation followed by sequencing (ChIP-seq) datasets of six histone modification marks (H3K27ac, H3K27me3, H3K6me3, H3K4me1, H3K4me3, H3K9me3) across 833 biosamples from the EpiMap repository at https://epigenome.wustl.edu/epimap/data/observed. These files were processed using macs2 ^84^ program, using -bdgpeakcall with default settings to call peaks. To determine association of peak breadth with variably expressed transcription factors (VETFs) (as described in ^29^) across all histone marks, we first used BEDTOOLs intersect ^85^ program to define peaks which overlapped with all protein-coding genes. For each biosample-histone modification combination, we binned peaks by their breadth into sets of 100 and evaluated the enrichment of VETFs (*n =* 634) across cumulatively decreasing peak breadth. All genes with no assigned peak were placed into the last bin. Fisher’s two-tailed test was used to determine enrichment. This process was repeated for a set of disease-associated long non-coding RNAs (lncRNAs) downloaded and combined from three independent databases (LncTard ^61^, LncRNADisease ^86^, MNDR ^87^). To determine the association of peak breadth with disease-associated lncRNAs, histone peaks overlapping with lncRNAs were binned into sets of 200 and enrichment was calculated.

### Source and processing of H3K27me3 ChIP-seq datasets

#### EpiMap human epigenomes (GRCh37/hg19)

We analyzed all observed and imputed ChIP-seq tracks of H3K27me3 across 833 human biosamples, grouped into 33 tissue categories, from the EpiMap repository at https://epigenome.wustl.edu/epimap/data/. These files were processed via macs2^84^ program, using -bdgbroadcall with default settings to call broad peaks. The median signal intensity for H3K27me3 peaks called across all 833 EpiMap^25^ biosamples was 0 (min) to 7.8 (max). Consolidating all peaks into one merged human H3K27me3 file, we found that the genome-wide coverage was 1,284,263,525 bases equating to 47.03% of the human genome (GRCh37/ hg19). We used liftOver to map all regions to GRCh38/hg38.

#### Roadmap human epigenomes

We analyzed all observed ChIP-seq tracks of H3K27me3 across 111 human biosamples, grouped from the Roadmap^88^ database at http://compbio.mit.edu/roadmap. These files were processed using macs2^84^ program, using -bdgbroadcall with default settings to call broad peaks. To facilitate comparison with the human EpiMap data, we rescaled the signal intensity to match the range of the EpiMap observed and imputed tracks. The rescaled signal intensity range used for human Roadmap H3K27me3 data upstream of macs2 peak-calling was rescaled to 0 (min) to 7.8 (max).

#### ENCODE human epigenomes

We analyzed all observed ChIP-seq tracks of H3K27me3 across X human biosamples from the ENCODE ^40^ database at https://www.encodeproject.org/. These files were processed using macs2 program, using -bdgbroadcall with default settings to call broad peaks. To facilitate comparison with the human EpiMap data, we rescaled the signal intensity to match the range of the EpiMap observed and imputed tracks. The rescaled signal intensity range used for human ENCODE H3K27me3 data upstream of macs2 peak-calling was rescaled to 0 (min) to 7.8 (max).

#### ENCODE mouse epigenomes

We analyzed all observed ChIP-seq tracks of H3K27me3 across 181 mouse biosamples, grouped into 31 tissue categories, from the ENCODE database at https://www.encodeproject.org/. These files were processed using macs2 program, using - bdgbroadcall with default settings to call broad peaks. We rescaled the signal intensity to match the range of the EpiMap observed and imputed tracks. The rescaled signal intensity range used for mouse ENCODE H3K27me3 data upstream of macs2 peak-calling was 0 (min) to 7.8 (max). Consolidating all peaks into one merged mouse H3K27me3 file, we found that the genome-wide coverage was 2,262,401,620 bases equating to 82.85% of the mouse genome (mm10). We note that the difference in H3K27me3 mark coverage between the human (47.03%) and mouse (82%) genomes is due to technical differences in ChIP-seq read depth and data-processing between the primary data sources used and likely not due to species-specific biological differences.

### Broad and consistent H3K27me3 domains mark cell-type specific regulatory loci

Given that facultative heterochromatin (fHC) formation is regulated by polycomb group (PcG) proteins ^24^, we use H3K27me3 marks as a proxy to demarcate patterns of fHC across the genome in this study. We aligned H3K27me3 ChIP-seq tracks across 833 EpiMap biosamples (i.e. cross-cell-type fHC alignment) and visualized peak (i) breadth and (ii) conservation over representative genomic loci (*GAPDH*, *MYH2*, *MYOD1* and *H19*). In each biosample, we categorized individual H3K27me3 peaks based on their breadth. A peak/domain was labelled as “broad” if its breadth ranked within the top 5% in that specific biosample, while all other peaks were classified as “non-broad”.

To evaluate the association between broad and conserved H3K27me3 domains (as measured by cellular constraint) with key developmental or cell-type-specific regulatory genes, we generated cellular constraint scores (CCS) for all NCBI RefSeq protein-coding genes and assessed the enrichment of representative positive gene-sets (**Table S1**). Genes were then ranked and binned based on their CCS, where each rank bin included 10% of all RefSeq genes evaluated with a CCS (*n =* 23,081). First, the proportion of genes involved in gene ontologies (GO) related to developmental, gastrulation, signaling and disease biological processes were evaluated across cumulatively decreasing CCS bins. Second, the enrichment of VETFs (*n =* 634) and TRIAGE priority genes (*n =* 1,359) from Shim *et al.* (2020) ^29^ were evaluated across cumulatively decreasing CCS bins using Fisher’s one-tailed test.

### Generating human cellular constraint scores at single-base resolution

(Supplementary Figures 2A & 2B)

For each base in the human genome we quantified the (i) breadth of overlapping H3K27me3 domains and (ii) conservation of H3K27me3 repression across 833 diverse biosamples in the EpiMap repository^25^. Within each biosample *j*, all H3K27me3 domain breadth values for peaks called *B_j_*={*b*_1*,j*_, *b*_2*,j*_,*b*_3*,j*_…) were normalized to yield domain breadth weightings (*w*) between 0 and 1, 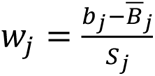 where 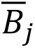 and *S_j_* are the sample mean and the standard deviation of H3K27me3 domain breadths in biosample *j*, respectively. The cellular constraint score (CCS) for each base is simply the sum of overlapping H3K27me3 domain breadth weightings across all 833 EpiMap biosamples. For each base in the human genome (*P*), the sum of the domain breadth weightings (*V_p_*) are calculated as follows.

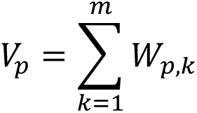

Finally, these were re-scaled into a range from 0 (lowest) to 1 (highest) to derive the cellular constraint score (*C_p_*) for each base, where *V_max_* and *V_min_* are the maximum and minimum sums of domain weightings across all bases.

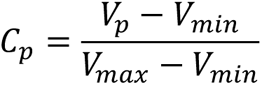

The result of this analysis quantifies a single CCS value > 0 for ∼1.3 billion bases (41.4%) of the human genome, which accumulatively forms a cellular constraint reference metric used for downstream analyses (GRCh37/hg19 and GRCh38/hg38 versions available). The remaining 58.6% of the genome displayed no detectable H3K27me3 signal in any of the EpiMap biosamples and were assigned a value of 0. We provide UCSC track sets for genome-wide human CCS for download at (https://genome.ucsc.edu/s/phycochow/hg19).

We also provide access to all CCS data files for reviewers here: https://cloud.rdm.uq.edu.au/index.php/s/7NzWP6fppAFErYe (Password: pLYHbWNbG7)

### Defining significantly cell-constrained regions

We defined significantly cell-constrained regions of the human genome by calculating the inflection point of the interpolated CCS curve of ∼1.3 billion bases using the ‘inflection’ package in R (v1.3.5) ^89^. Overall, we defined 43,245,600 significantly cell-constrained bases accounting for ∼1.4% of the human genome. Of the ∼43 million significantly cell-constrained bases, we found approximately equal contribution of coding (∼19 million) versus non-coding (∼21 million) bases. For each input dataset used in this study, the BEDTOOLs intersect program ^85^ was used to identify input regions overlapping with the CCS reference metric to calculate an average CCS for each input region.

### General characterization of cell-constrained regions

To assess the genomic distribution of significantly cell-constrained domains, we compared their H3K27me3 domain breadths and CCS values in relation to distance from the nearest RefSeq protein-coding gene. Cell-constrained domains were grouped into different categories on this basis (NCC= non-cell-constrained, CC= cell-constrained, G= gene-overlapping cell-constrained, I= intergenic cell-constrained). A domain was classed as ‘gene-overlapping’ if it overlapped entirely with a RefSeq protein-coding gene using the BEDTOOLs intersect program ^85^. All other domains were classed as ‘intergenic’. Subsets of intergenic categories were defined using lenient (I= +/- 0kb around RefSeq gene body), moderate (10kb-I= +/-10kb), intermediate (30kb-I= +/-30kb) and stringent (50kb-I= +/-50kb) cut-offs based on varying distances +/- from the chr start and end position of the RefSeq gene-body.

To evaluate whether there was sufficient cellular diversity available in available epigenomic databases to accurately determine cellular constraint in humans, we tested the correlation between single-base cellular constraint scores from 833 EpiMap biomsaples versus 111 Roadmap biosamples using Spearman’s rank correlation test (Spearman’s *r*= 0.9208, *p*=<0.0001).

### Generating mouse cellular constraint scores at single-base resolution

For each base in the mouse genome, we quantified the (i) breadth of overlapping H3K27me3 domains and (ii) conservation of H3K27me3 repression across 181 diverse biosamples in the ENCODE database ^40^ using the same method as described for human CCS calculation. The result of this analysis quantifies a single CCS value for ∼2.3 billion bases in the mouse genome, which accumulatively forms a cellular constraint reference metric used for downstream analyses (GRCm38/mm10 version available). We provide track sets for genome-wide mouse CCS for download at (link to UCSC tracks; REFF). We defined significantly cell-constrained regions of the mouse genome by calculating the inflection point of the interpolated CCS curve of 2.3 billion bases using the ‘inflection’ package in R (v1.3.5) ^89^. Overall, we defined 40,894,665 significantly cell-constrained bases accounting for ∼1.5% of the mouse genome. We provide UCSC track sets for genome-wide mouse CCS for download at (https://genome.ucsc.edu/s/phycochow/mm10).

### Correlation between mouse and human cellular constraint scores

To retrieve syntenic regions between human (GRCh37/hg19) and mouse (GRCm38/mm10) genomes, we used the SynBuilder application with the minimum resolution set at 150,000bp on the Synteny portal at https://bioinfo.konkuk.ac.kr/synteny_portal. We filtered this file to keep only genomics regions that mapped in both directions (i.e. in both the hg19 to mm10 file and the mm10 to hg19 file) termed consensus regions. These consensus regions were then divided into 100bp blocks and used for correlation analysis.

To evaluate the conservation of broad and conserved H3K27me3 patterns across different species, we tested the correlation between cellular constraint scores for 100bp syntenic regions calculated in the human versus the mouse genomes using Spearman’s rank correlation test (Spearmans’s *r*= 0.593, *p*=<0.0001).

### Permutation overlap analysis of evolutionary signals

To statistically evaluate the associations between cell-constrained regions and signals of evolutionary conservation and adaptation, we performed permutation tests using the regioneR ^43^ package. To determine the significance of the overlap between cell-constrained bases and selected evolutionary signals (data sources listed below), we used the ‘overlapPermTest’ function with ‘randomizeRegions’ performed with either 1000 or 10,000 permutations.

*PhastCons conserved elements*: PhastCons 100-way vertebrate conservation scores were downloaded at http://hgdownload.soe.ucsc.edu/goldenPath/hg19/phastCons100way. PhastCons elements were filtered using a score cut-off ≥ 0.9 to keep only highly sequence conserved bases for analysis (*n*= 12.4×10^7^ bases). *Human specific structural variants (hsSVs):* hsSVs (*n=* 17,789 variants) were downloaded directly from supplementary material provided in Keogh *et al.* (2023) ^50^. *Human accelerated regions (HARs):* High-confidence zooHARs (*n=* 312) were downloaded directly from the supplementary material provided in Keogh *et al.* (2023) ^50^. *Archaic adaptive introgression (AI):* Regions of adaptive introgression (*n*=862) were downloaded combined from Gittelman *et al.* (2016) ^46^, Gower *et al.* (2021) ^90^, Racimo *et al.* (2017) ^45^, and Setter *et al.* (2020) ^91^.

### Genealogical estimation of variant age

Genealogical Estimations of Variant Age (GEVA) were retrieved from the atlas of variant age at https://human.genome.dating/. Variants were filtered for common variants only with a minor allele frequency (MAF) greater than 0.01. Cellular constraint scores were calculated for each variant. The combined mutational age estimates from the TGP and SGDP databases was preferentially used where possible. We defined significantly cell-constrained variants using the inflection point of the interpolated CCS curve. Variants were binned into 100 discrete bins based on their reference allele frequency. The average allele age within each of these bins was calculated for cell-constrained and non-constrained regions. To determine statistical significance at each MAF bin, we performed bootstrapping of non-constrained variants to establish an expected distribution from which we derived a Z-score for the significantly cell-constrained regions.

### Long non-coding RNA enrichment analyses

We downloaded lists of curated long non-coding RNAs (lncRNAs) from various published databases including evolutionarily conserved lncRNAs ^30^, SyntDB ^53^, developmentally dynamic lncRNAs ^30^, LNCipedia ^92^, LncTard ^61^, LncRNADisease ^86^, MNDR ^87^, and yin-yang lncRNAs ^93^. Cellular constraint scores were generated for all LNCipedia transcripts. We defined 5,723 significantly cell-constrained lncRNAs using the inflection point of the interpolated CCS curve. LncRNA transcripts were matched across databases using Ensembl IDs. The enrichment of cell-constrained lncRNAs within the curated lncRNA sets was calculated using Fisher’s two-tailed test.

For binned analyses, all lncRNAs were ranked by CCS and binned into a percentile bin. At each rank bin, we quantified the enrichment of a given curated lncRNA set using a Fisher’s two-tailed test. This represents how significantly a curated lncRNA set is enriched among lncRNAs with CCSs above a certain rank (i.e. in the top x% of all lncRNAs ranked by CCS) compared to the rest using a sliding window.

### Analysis of CAGE-sequencing data from time-course of hiPSC cardiac differentiation

Bulk CAGE-sequencing data capturing four time-points (Days 0, 2, 5 and 20) of human induced pluripotent stem cell (hiPSC) cardiac differentiation was generated in-house. The CAGE-seq data was analyzed using the CAGEr pipeline as previously described ^94^. In this analysis, we aimed to demonstrate how cellular constraint annotation could be integrated with complementary transcriptional data as a weighting metric to inform cell-type-specific function and accelerate novel gene discovery pipelines (**see Supplementary text**).

For each time-point, we generated CCSs (CCS_t_) for all CAGE transcription start sites (TSSs) and used it to weight the normalized transcript abundance (T_t_) (i.e. expression) to generate a CCS-weighted expression values (WT _t_) for each TSS, as follows:

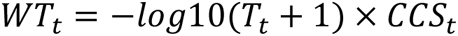

The CCS-weighted expression metric accounts for a genomic element’s tendency to be epigenetically safeguarded by broad H3K27me3 across many cell-types, as well as its observed transcriptional abundance within a given cell- or tissue-type. In this analysis, CCS-weighted expression values were used to rank TSSs at each differentiation time-point and prioritize transcripts with cell-type specific regulatory function for downstream experimental validation.

### Candidate transcript selection for downstream validation

Candidate transcripts for biological validation were selected using the following criteria (as outlined in **fig. S4C**): **Phase I: Step 1.** Identify CAGE transcripts that are novel or unannotated within each population. **Step 2.** Shortlist transcripts ranked within the top 5% of CCS-weighted expression values. **Step 3.** Select transcripts with no known role published in existing literature. **Step 4.** Select transcripts that are either nearby a transcription factor or be completely intergenic with very high individual CCS ranking. At this stage we shortlisted four candidate lncRNAs for Phase II: *RP11-175E9.1, AL662890.1, lnc-NKX2-5-1:1, RP11-89K21.2*. **Phase II: Step 1.**

Generate CRISPRi loss-of-function hiPSC lines targeting the four candidate transcripts. **Step 2.** Run experiments to quality control and measure the efficiency of transcript knockdown. **Step 3.** Perform functional assays and phenotyping on any candidates that pass all check.

| Candidate transcript | Chr location (hg38) | Population of interest | Nearby TF | Efficient CRISPRi K/D |
| --- | --- | --- | --- | --- |
| RP11-175E9.1 | chr8:23563809-23566144 | Day 5 & 20 hiPSC cardiac differentiation | Directly upstream of NKX2-6 | Yes |
| AL662890.1 | chr6:28743281-28744400 | Day 2 hiPSC cardiac differentiation | >100kb away | No |
| lnc-NKX2-5-1:1 | chr5:172669563-172671964 | Day 20 hiPSC cardiac differentiation | ~ 7.5kb upstream of NKX2-5 | No |
| RP11-89K21.1 | chr2:45161052-45163229 | Day 2 hiPSC cardiac differentiation | ~ 3kb upstream of SIX3 | No |

Based on this selection criteria, *RP11-175E9.1* was the only candidate that passed all checks and was selected for downstream functional screening and phenotype validation experiments.

### Maintenance and Culture of Human iPSC Lines

All human pluripotent stem cell studies were carried out in accordance with consent from the University of Queensland’s Institutional Human Research Ethics approval (HREC#: 2015001434). WTC CRISPRi GCaMP hiPSCs (Karyotype: 46, XY; RRID: CVCL_VM38) were generated using a previously described protocol^95^ and were generously provided by M. Mandegar and B. Conklin (UCSF, Gladstone Institute). WTC CRISPRi RP11-175E9.1-g1 hiPSCs were generated in this study (see below). All hiPSCs were maintained on Vitronectin XF (Stem Cell Technologies, Cat.#07180) coated plates, in mTeSR+ media with supplement (Stem Cell Technologies, Cat.#05850) at 37° C with 5% CO2.

Using a modified monolayer-based hiPSC differentiation protocol based on previous reports^96^, we differentiate hiPSCs into multiple cell lineages to derive a mixture of heterogenous cell populations. On day -1 of differentiation, hPSCs were dissociated using 0.5% EDTA, plated into vitronectin coated plates at a density of 1.8 x 10^5^ cells/cm^2^ (24 well-plate) or 2 x 10^4^ cells/cm^2^ (T25 flask), and cultured overnight in mTeSR media (Stem Cell Technologies). Differentiation was induced on day 0 by first washing with PBS, then changing the culture media to RPMI (ThermoFisher, Cat.#11875119) containing 3μM CHIR99021 (Stem Cell Technologies, Cat.#72054), 500μg/mL BSA (Sigma Aldrich, Cat.#A9418), and 213μg/mL ascorbic acid (Sigma Aldrich, Cat.#A8960). After 3 days of culture, the media was replaced with RPMI containing 500μg/mL BSA, and 213μg/mL ascorbic acid. On day 5, the media was exchanged for RPMI containing 500μg/mL BSA, and 213μg/mL ascorbic acid without supplemental cytokines. Cells were cultured in this media till harvest on Day 6. The differentiation cultures were maintained at 37°C in a 5% CO_2_ incubator.

### Generation of WTC CRISPRi *RP11-175E9.1*-g1 hiPSCs

Three separate guide RNAs (gRNAs) targeting the CAGE-defined transcriptional start site of the human RP11-175E9.1 sequence, p@ENST00000523874, were designed and cloned into the pQM-u6g-CNKB doxycycline-inducible construct and transfected into WTC CRISPRi GCaMP hiPSCs using the Neon transfection system (Invitrogen, Cat.#MPK1096). For electroporation, 0.5μg DNA and 1×105 dissociated hiPSCs were mixed in 10μL resuspension buffer R (Invitrogen, Cat.#MPK1096). Electroporation parameters were as follows: pulse voltage, 1300V; pulse width, 30ms; and pulse number, 1. Cells were then plated in Vitronectin XF (Stem Cell Technologies, Cat.#07180) coated plates in mTeSR media (Stem Cell Technologies, Cat.#05850) supplemented with 10μM of Y-27632 (Stem Cell Technologies, Cat.#72308). Stable clones were selected using successive rounds of re-plating with blasticidine at 10μg/ml (Sigma, Cat.#15205). Populations were tested for knockdown efficiency by qPCR following doxycycline addition at 1 μg/ml (Sigma, Cat.#D9891) continuously from day 0 of cardiac-directed differentiation. Of all the cell lines generated, the WTC CRISPRi RP11-175E9.1-g1 line displayed high knockdown efficiency and therefore was chosen for downstream functional testing.

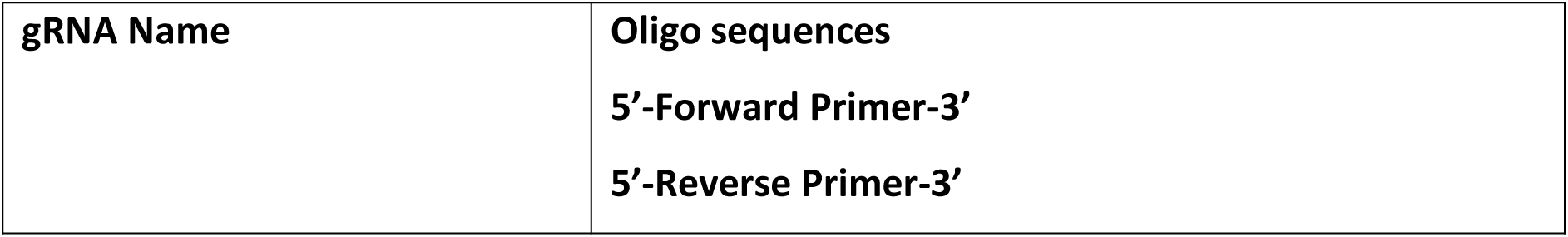
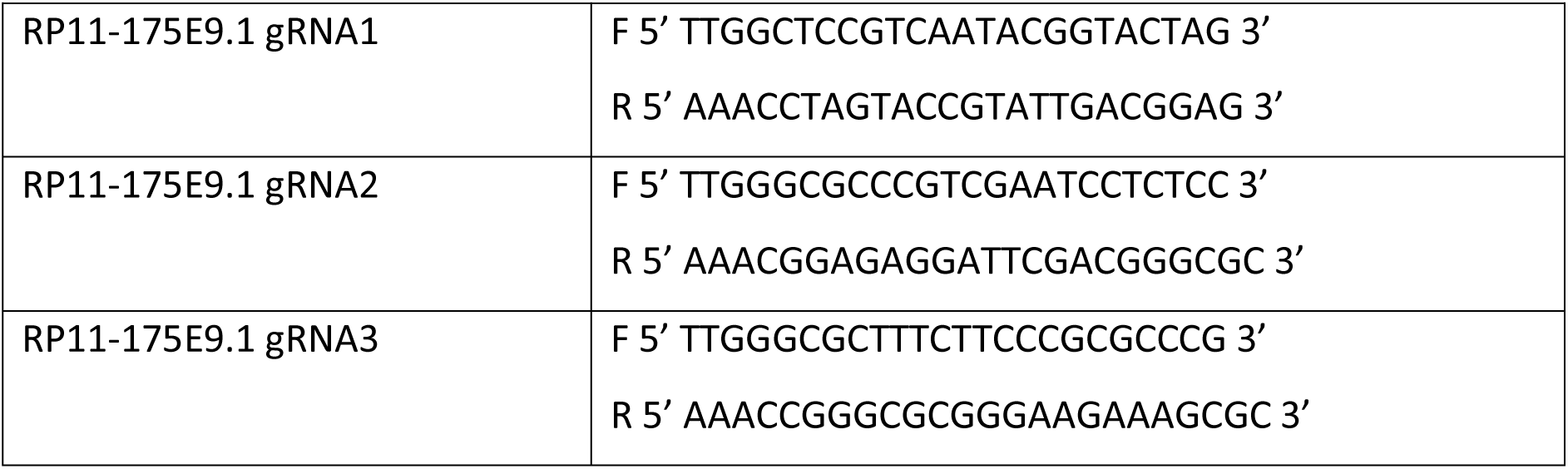

### Quantitative RT-PCR

For quantitative RT-PCR, total RNA was isolated using the RNeasy Mini kit (Qiagen, Cat.#74106). First-strand cDNA synthesis was generated using the Superscript III First Strand Synthesis System (ThermoFisher, Cat.#18080051). Quantitative RT-PCR was performed using SYBR Green PCR Master Mix (ThermoFisher, Cat.#4312704) on a ViiA 7 Real-Time PCR System (Applied Biosystems). The copy number for each transcript is expressed relative to that of housekeeping gene HPRT1. Samples were run in biological triplicate. FC was calculated on a gene by gene basis as gene expression divided by control gene expression. The following are qRT-PCR primers utilized in this study:

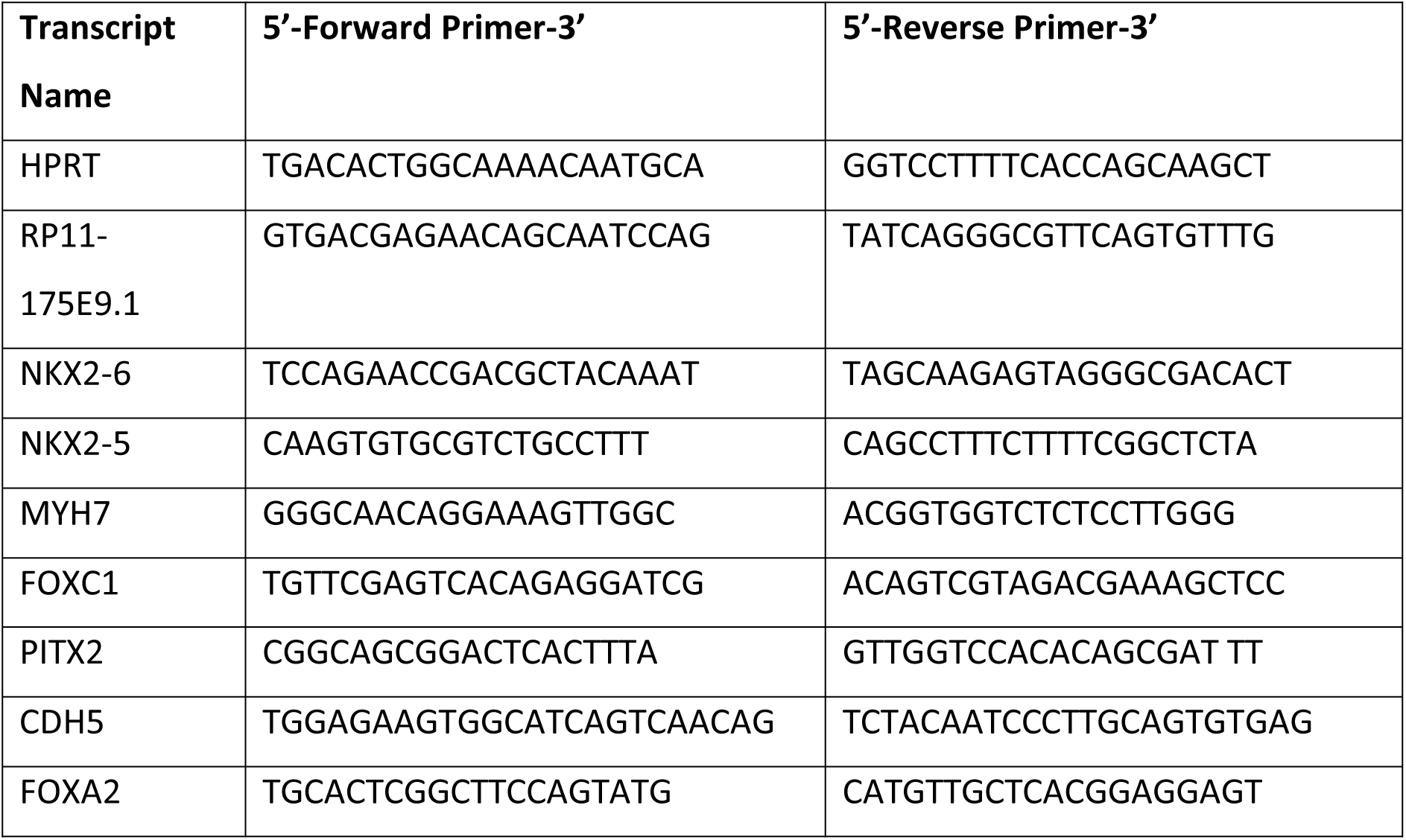
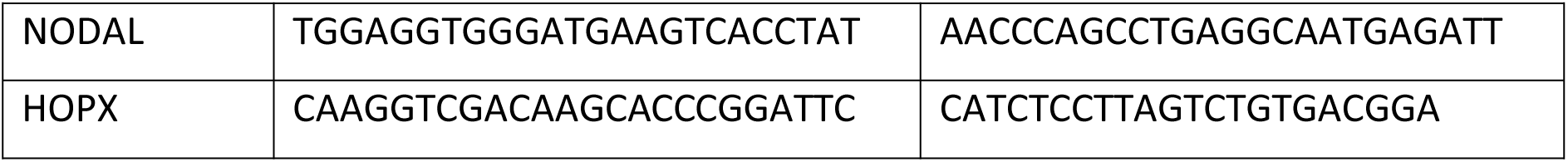

### Flow cytometry analysis

On day 15 of cardiac differentiation cells were fixed with 4% paraformaldehyde (Sigma, Cat.#158127) and permeabilized in 0.75% saponin (Sigma, Cat.#S7900). Fixed cells were labelled for flow cytometry using cardiac troponin T (cTnT) (ThermoFisher Scientific, Cat.#MA5-12960; RRID: AB_2206583) and corresponding isotype control. Cells were analysed using a BD FACSCANTO II (BD Biosciences) with FACSDiva software (BD Biosciences). Data analysis was performed using FlowJo (Tree Star).

### Single cell RNA-Sequencing

#### Sample handling and library preparation

Cells were harvested using 0.5% EDTA + Trypsin (1:10) solution and neutralised using RPMI with FBS (1:1). Samples relevant to this project were multiplexed as part of a larger experimental run using a mixture of internal and external barcode labelling. Cells from the experimental conditions relevant to this project were stained individually with their assigned cell-hashing antibody (TotalSeq-A anti-human cell-hashing antibodies). All samples were pooled and sorted for viability on a Beckman Coulter MoFlo Astrios EQ Cell Sorter with a ZombieDye stain (BioLegend). For each condition, 5×10^5^ live cells were collected and 2×10^4^ were run on the 10X Chromium instrument for single-cell capture, in accordance with the manufacturer’s protocol. Following the cell-hashing protocol^97^, the transcriptome and antibody barcode libraries were prepared separately. The transcriptome libraries were sequenced on the Illumina NovaSeq 6000 and the cell-hashing libraries on the Illumina NextSeq 550Dx.

#### Demultiplexing and quality control

Barcoding reads from separately amplified and sequenced barcode libraries were used to sample barcodes to cells. Cells without both transcriptome and sample barcoding reads were removed. For each barcode sequencing library, the ‘HTODemux’ function in the Seurat R package (v3.0) was used to determine the dominant sample barcode for each cell and annotate negative and doublet cells based on their sample barcode reads alone. Three transcriptome-based doublet detection methods in the scds R package (v1.6.0) were used to further assign doublet annotations to each cell, and cells labelled as doublets by at least three methods were removed. Transcriptome-based cell filtering as part of the Seurat pipeline ^98^ removed cells with fewer than 1500 and greater than 6000 detected genes; fewer than 4000 and greater than 37,500 total read counts; mitochondrial reads accounting for greater than 20% of total reads; or cells with greater than 40% gene expression from ribosomal genes. Following filtering, sample barcodes were assigned to the remaining cells based on the barcode with the highest expression in each cell.

#### Dataset Integration

Sample libraries were generated and sequenced in separate experimental batches to generate two scRNA-seq datasets. Data integration on both scRNA-seq batches was performed implementing the Reference Principal Component Integration (RPCI) method using the RISC R package (v1.0) ^99^. RPCI utilizes the gene eigenvectors from a reference dataset to establish a global frame for integration of scRNA-seq data from multiple batches or studies. Specifically, we combined individual datasets using the scMultiIntegrate function of the RISC package and used the output corrected gene expression values for all downstream analyses in Seurat (v4.0.5) ^98^.

### Downstream analysis of scRNA-seq data

Normalisation, UMAP dimensionality reduction, and clustering of the data was done following the standard Seurat (v4.0.5) pipeline ^98^. The clustering resolution used was 0.2, and we assigned cell type labels by interrogating marker gene expression in each cluster using the Seurat ‘FindAllMarkers’ function (min.pct = 0.25, logfc.threshold = 0.25).

### Promoter capture Hi-C data analysis

(Supplementary text)

We used publicly available promoter capture Hi-C (PCHi-C) data from both hiPSCs and hiPSC-derived cardiomyocytes (hiPSC-CMs) ^100^. Using the BEDTOOLs ^85^ intersect program we identified promoter regions interacting with the *RP11-175E9.1* sequence. From this we identified the following *RP11-175E9.1*-promoter interactions in hiPSCs and hiPSC-CMs:

| Promoter interactions in hiPSCs | Promoter interactions in hiPSC-CMs |
| --- | --- |
| LOXL2 | R3HCC1 |
| NKX2-6 | NKX2-6 |
| ENTPD4 | ENTPD4 |
| SLC25A37 | SLC25A37 |
| NKX3-1 |  |

### Analysis of GTEX eQTL data

All significant *cis*-eQTL gene pairs were downloaded for 49 tissue types from publicly available data released by GTEx consortium at https://www.gtexportal.org/home/datasets. CCSs were calculated for all eQTLs across all tissues. All variants without a CCS were excluded. We defined significantly cell-constrained eQTLs using the inflection point of the interpolated CCS curve. A Wilcoxon’s signed rank test was used to compare the regression slope and significance of the eQTL-gene associations between cell-constrained eQTLs and non-constrained eQTLs. Each eQTL-gene association was treated individually, including those that featured the same eQTL but were associated to a different gene. Wilcoxon’s signed rank test was used to compare the minor allele frequency between cell-constrained and non-constrained variants, with duplicate eQTLs removed for this analysis.

### Enrichment of fine-mapped causal variants from Kanai *et al.* (2021)

CCS scores were calculated for each of the 908,787 variant-trait pairs for which fine-mapping had been performed by Kanai *et al.* (2021) ^69^. Of these 2,108 had a high posterior inclusion probablility (PIP) of being the causal variant (PIP > 0.9) and an assigned CCS score. We defined 48,299 significantly cell-constrained variants using the inflection point of the interpolated CCS curve. Enrichment for causal variants among cell-constrained variants was calculated using fisher’s exact test. The proportion of variants that were cell-constrained across different PIP values was calculated for 100 bins between PIP 0 and PIP 1.

### Analysis of ClinVar pathogenic variants

All variants for which disease or non-pathogenic data was reported to the ClinVar database were downloaded (https://www.ncbi.nlm.nih.gov/clinvar/). Variants were combined from the GRCh37 and GRCh38 assemblies using the unique variant identifier to ensure there were no duplicates. Cellular constraint scores were calculated for each variant and variants without a CCS (CCS ≤ 0) were excluded from the analysis. A total of 40,294 variants out of 560,694 were defined as significantly cell-constrained based on the inflection point of the interpolated CCS curve. Their genomic location was annotated per the information provided by ClinVar, denoting variants as coding or non-coding (genomic). Variants classified as pathogenic or likely pathogenic in the database were defined as pathogenic for this analysis. A fisher’s exact test was used to compare the frequency of cell-constrained variants among pathogenic variants compared to non-constrained variants. Odds ratio was calculated to determine if variants were over-represented or under-represented. Variants were stratified by their genomic location into three categories, all, coding and genomic. Enrichment was calculated across 30 equally spaced CCS bins for the three variant groups. This analysis was repeated on variants which had a disease associated with them with ‘congenital’ in the disease trait name.

### SBayesRC heritability enrichment analysis of 28 UKBB complex traits and diseases

We selected 28 traits from UKBiobank with relatively large sample size and pair-wise phenotypic correlation |r| < 0.3. The phenotype out the range of mean +/- 7SD were filtered and then was rank-based inverse-normal transformed within each ancestry and sex group. To validate the prediction performance, ten-fold cross-validation were performed in the unrelated EUR samples (n=341,809). Ten equal-sized disjoint samples were partitioned, and in each fold, one subsample was used as the validation, while the other nine subsamples were used as training. For the training summary statistics, linear regression (quantitative trait) or logistic regression (binary traits) were run in PLINK2.0.

We generated cellular constraint scores for all SNPs which were used in the continuous CCS annotation model. Using the inflection point of the interpolated CCS curve we defined a subset of significantly cell-constrained SNPs which were assigned 1 and all other SNPs 0, representing the binary CCS annotation model. We input the summary data from training data, LD matrix downloaded from SBayesRC data repository, and with or without cellular constraint annotation to SBayesRC R packages (version 0.2.0) ^101^. Polygenic risk scores (PGS) were calculated in validation set by PLINK2.0, and the prediction R^2^ was obtained from linear regression of phenotypes on the PGS for quantitative traits, and McFadden’s pseudo-R^2^ from logistic regression for binary traits. The final R^2^ was calculated from the R^2^ from the full model (PGS + sex + age + 10 PCs) - the null model (sex + age + 10 PCs). The relative prediction accuracy was calculated 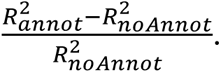 We also obtained the prediction accuracy from SNPs annotated with CADD ^75^ and 29-way mammalian constraint ^76^. To evaluate if cellular constraint was complementary to commonly used genomic annotations, we calculated the prediction accuracy with the annotation CADD + 29-way mammalian constraint + binary CCS, and compared to CADD + 29-way mammalian constraint model. We found with the binary CCS annotation, the prediction accuracy improved 0.3% averaged from all trait (Wilcox test, V = 385, p-value = 3.33e-06).

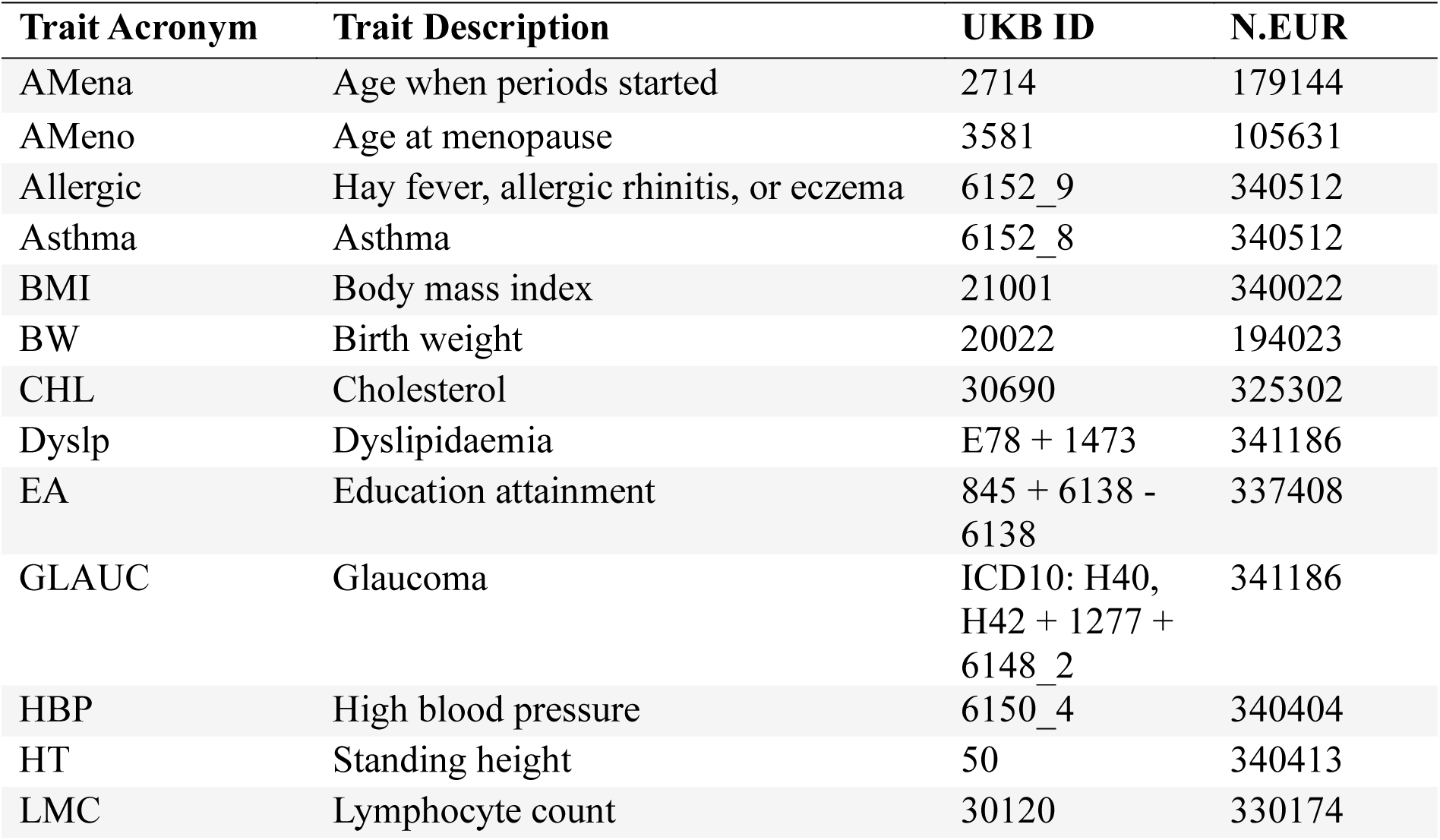
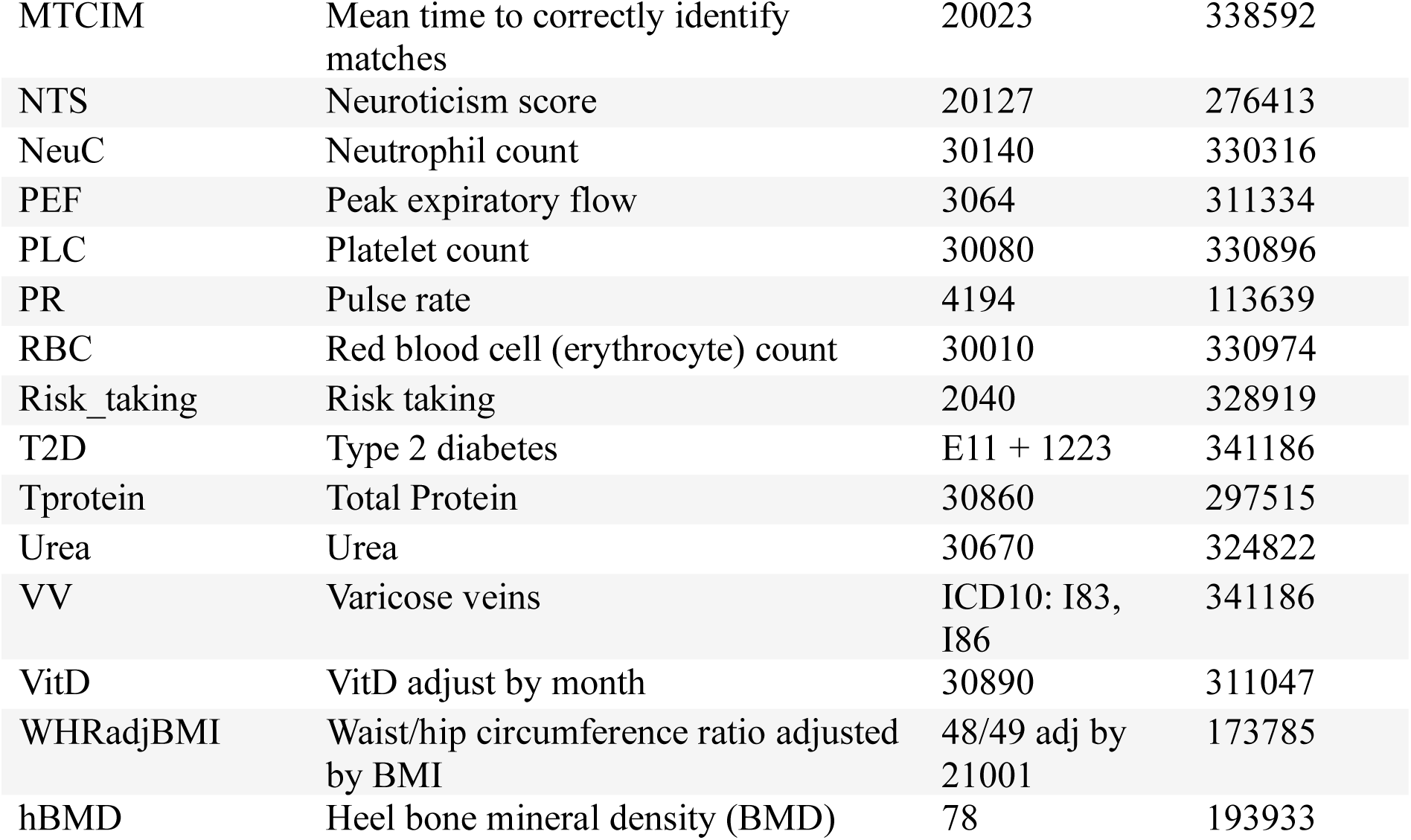
Table. Description of 28 UK Biobank complex traits and diseases analysed in this study.

### CARMA functionally informed fine-mapping analysis

The ‘CARMA’ R package (version 1.0) ^102^ was utilized to perform fine-mapping using the UK biobank GWAS summary statistics of EUR ancestry for 25 complex traits and diseases. For cancer-related traits, we obtained GWAS meta summary statistics of breast cancer ^79^ from EUR ancestry. LD matrix was obtained from 337k unrelated UKB individuals of EUR ancestry at https://alkesgroup.broadinstitute.org/UKBB_LD. For each 3Mb region harboring a significant locus of a certain trait, a statistical fine-mapping was conducted with three different annotation strategies; no functional annotation, binary cellular constraint annotation, and continuous cellular constraint annotation. Posterior inclusion probabilities (PIP) were summarized for each variant of tested regions. Outlier detection was switched off for the UKB data due to the consistent data resource between summary statistics and LD reference.

### Machine learning survival modelling in cancer cohorts

We generated CCS values for a list of variants (single-base and CNVs separately) and used the inflection point of the CCS curve to define 1,123 significantly cell-constrained variants which represented the ‘cell-constrained variant panel’. This was compared to an ‘expert curated variant panel’ of 525 variants used as a clinical gold standard for cancer genomic testing. Both variant panels were tested in their ability to add useful information to a kernel survival model trained on cancer patients from The Cancer Genome Atlas Program (TCGA) ^103^, by using them as a feature for a kernel function.

The survival model uses mutation features in a *Random Forest Kernel* (similar to a jaccard index metric) with tunable scaling and weighting at the output. Repeated 5-fold cross-validation was used as the test, for both variant panels defined. For each round of cross-validation, the kernel parameters were trained to minimize negative log-likelihood in the training data. The trained kernels were then used to estimate a survival curve for each subject in the testing set and the survival curves used to estimate 5-year survival. The model was assessed on the AUC of this 5-year survival classifier. For a full description of the machine-learning model and implementation on multimodal (including genomic) data, see Griffiths *et al.* (2024) ^78^.

Kernel survival modelling is a survival modelling technique that extends traditional survival modelling. Each subject has a unique hazard function, with estimation of the hazard function produced by a modified *Kaplan-Meier* estimator. The subject count in the numerator and denominator of the *Kaplan-Meier* estimator is replaced by a sum of a similarity measure between the subject for which the hazard function is being estimated, and the other subjects. The function that produces the similarity measure between pairs of subjects is known as a kernel function.

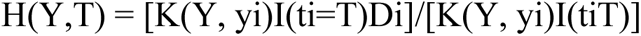

Where:

H is the hazard for person Y at time T

- [0:1], probability a person does not survive to a particular time point given they have survived up until that time point.
I is the indicator function

- 1 for true input, 0 for false input.
y and t are the set of training subjects

- A feature vector – such as a binary variable for 0 or 1 for presence or absence respectively of a genomic variant, e.g. TP53 mutation.
- The time of their event (e.g. 100 days after diagnosis).
D is a binary variable for each training subject

- 1 for a death and 0 for a censorship.

One way to think of the *Kaplan-Meier* estimator is in relation to the more general kernel model. In this way, the *Kaplan-Meier* estimator is a kernel estimator where the kernel function is strictly one.

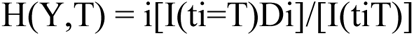

And in both the naive always-one case, and the full kernel case, it holds that for person Y at time T:

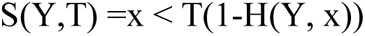

In summary, S(Y,T) is the standard *Kaplan-Mier* survival curve estimator. In effect, this output is the probability of survival (1 - H) at a specific time (x) for an individual (Y), given the reported features. The reported features in this study compared the performance of the expert curated variant panel against the cell-constrained variant panel.

### Drug approval prediction analysis

To investigate if the cellular constraint score of a drug’s gene target could be used as a predictor of drug approval we utilised the Pharmaprojects dataset published in King *et al* (2019) ^82^. This dataset contains indication-drug-gene trios and their approval status for clinical use. We integrated the CCS with the success probability defined by King et al (2019). Success probabilities with no outcome in the Pharmaprojects database were removed. Further, the strongest success probability per gene was retained to remove duplicate gene records, leaving 708 drug-gene-indication trios. We performed cumulative binning of genes, with the average CCS decreasing as the number of bins included increased. We calculated the success probability in each of these bins separately for approved and non-approved drugs. Further, we calculated the AUC for predicting approval status based on the success probability at each CCS bin.

## Acknowledgements

This work has been supported by grant funding to N.J.P. (ARC SR1101002, DP170101217, NHMRC APP1143163, and National Heart Foundation FLF106721).

## Author contributions

E.S. conceptualized the study, performed computational analyses and functional assays, interpreted results, and wrote the manuscript. D.M. performed allele age, GTEx, ClinVar, and drug prediction analyses. W.J.S. and C.S.Y.C. generated human and mouse CCS values. Y.S. performed analysis of AI regions. F.F.C. performed fine-mapping analysis. Z.Z. performed SBayesRC analysis. J.L. performed machine learning analysis. M.F., S.S., M.B., J.Z., B.L. contributed intellectually and provided feedback on the manuscript. N.J.P. conceptualized the study, supervised the project, raised funding, and edited the manuscript.

## Materials & Correspondence

Further information and requests for resources and reagents should be directed to and will be fulfilled by the corresponding author N.J.P.

## Data and materials availability

Data are made available for reviewers to access here: https://cloud.rdm.uq.edu.au/index.php/s/7NzWP6fppAFErYe (Password: pLYHbWNbG7). These data files will be made available for public access upon publication. All other data are available in the manuscript or the supplementary materials.

## Supplementary Figures

**Supplementary Figure 1.**
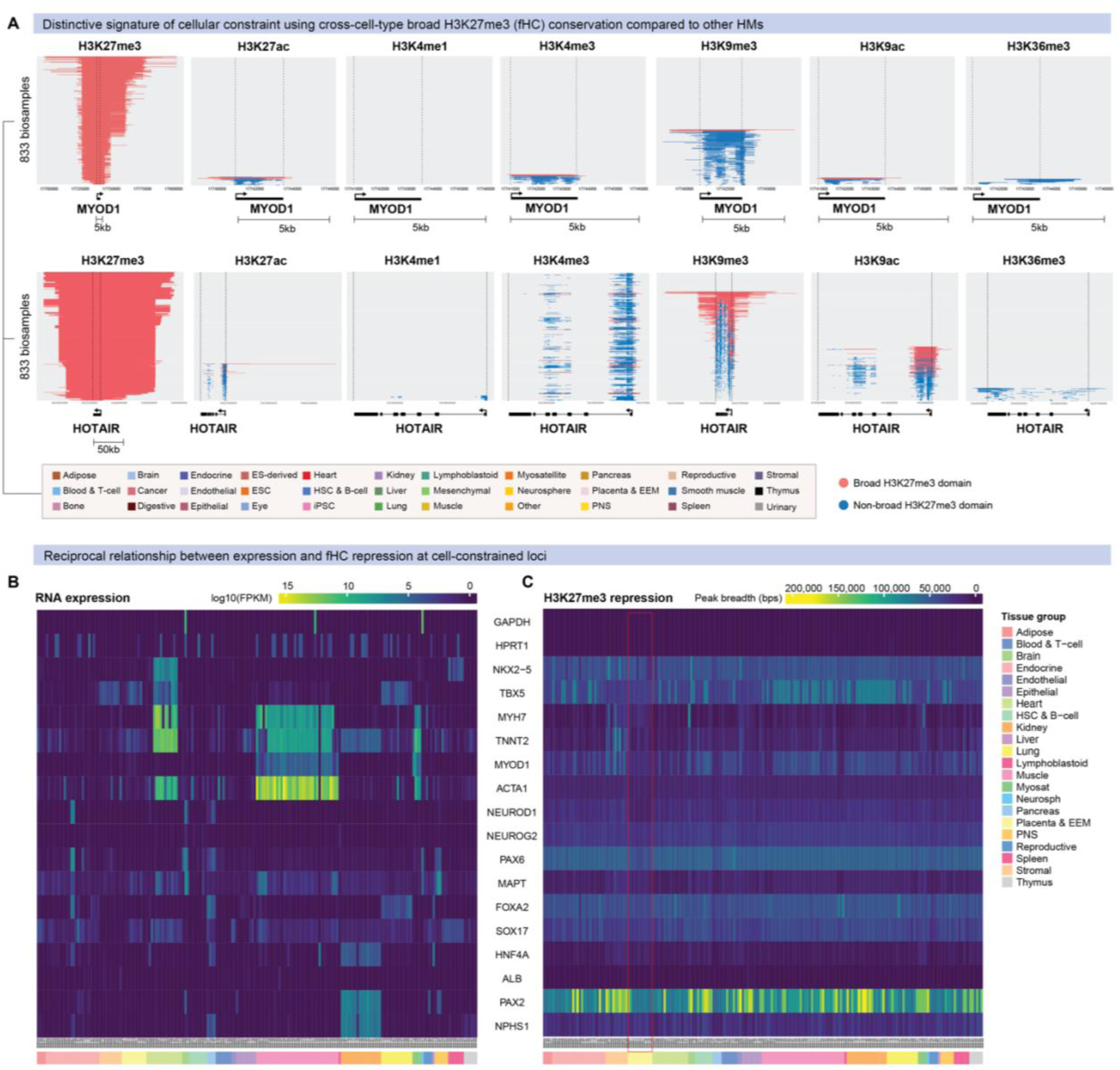
Cross-cell-type histone modification ChIP-seq alignment across diverse cell- and tissue-types. **(A)** Plots display aligned ChIP-seq peaks from 833 EpiMap biosamples, showing deposition of seven HMs over a locus stacked vertically and sorted on breadth. Domains ranked in the top 5% within each biosample are considered broad (red) and the remainder non-broad (blue). (**B)** Heatmaps displaying RNA expression (log10 FPKM) (left) and H3K27me3 repression (peak breadth) (right) of representative genes.

**Supplementary Figure 2.**
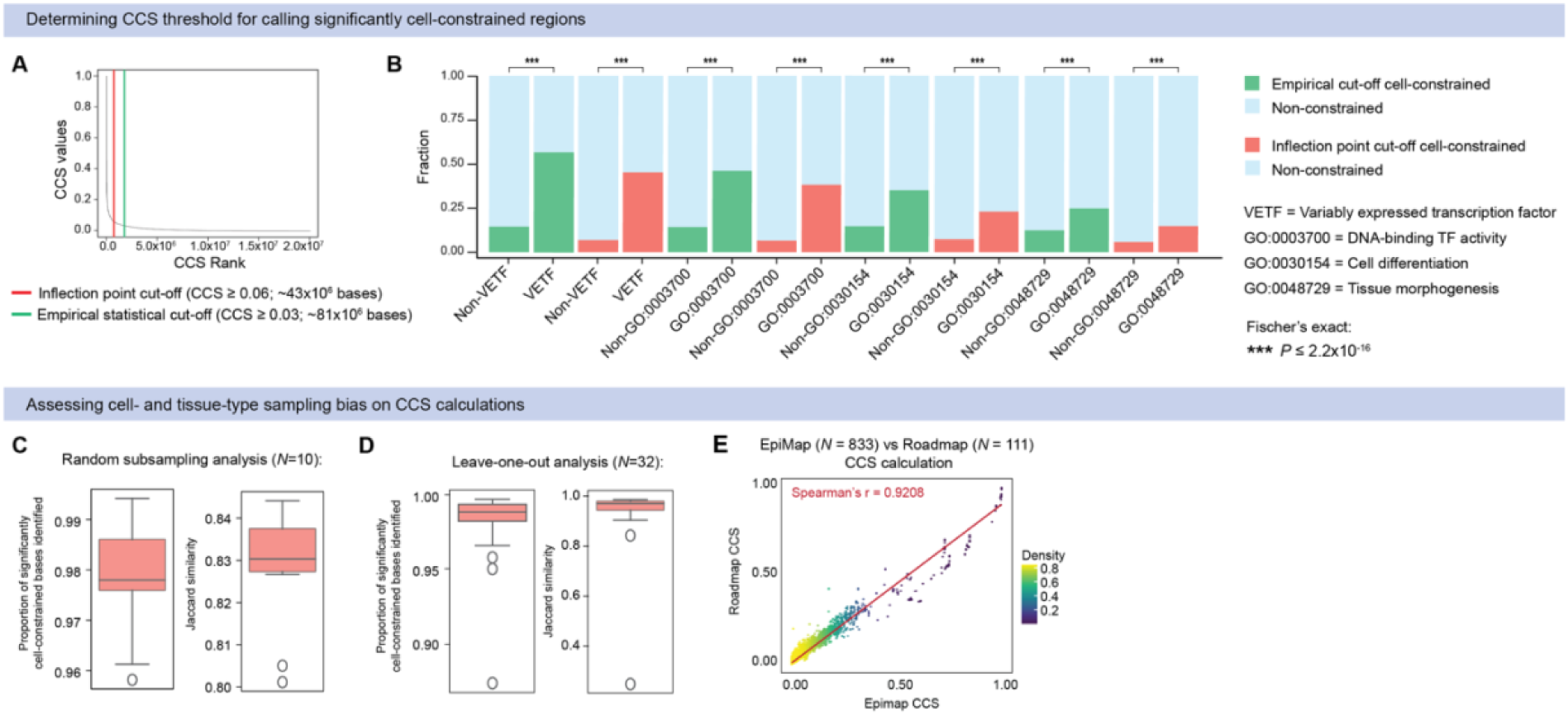
General characterization of human cellular constraint metric. **(A)** Distribution of single-base CCS values showing thresholds for calling significantly cell-constrained bases using the inflection point cut-off (CCS ≥ 0.06; ∼43×10^6^ bases) (red line) vs. an empirical statistical cut-off (CCS ≥ 0.03, ∼81×10^6^ bases) (green line). (**B**) Comparing the enrichment of representative gene-sets using the inflection point versus an empirical statistical cut-off to determine significantly cell-constrained bases (\*\*\**P≤*2.2×10^-16^, two-tailed Fisher’s exact test). (**C-E**) Evaluating cell- or tissue-type bias within biosamples effect on CCS calculations. (**C**) Bootstrapping analysis via random resampling of 833 biosamples (*N*= 10 permutations) showing proportion (left) and jaccard similarity index (right) of significantly cell-constrained bases identified. (**D**) Leave-one-out analysis via iterative removal of all biosamples from each of the 33 tissue-groups represented (*N*= 32 permutations) showing proportion (left) and jaccard similarity index (right) of significantly cell-constrained bases identified. (**E**) Correlation between cellular constraint scores calculated used epigenomes from EpiMap (*N=* 833 biosamples) versus Roadmap (*N=* 111 biosamples) (Spearman’s *r*= 0.9208, *p*=<0.0001).

**Supplementary Figure 3.**
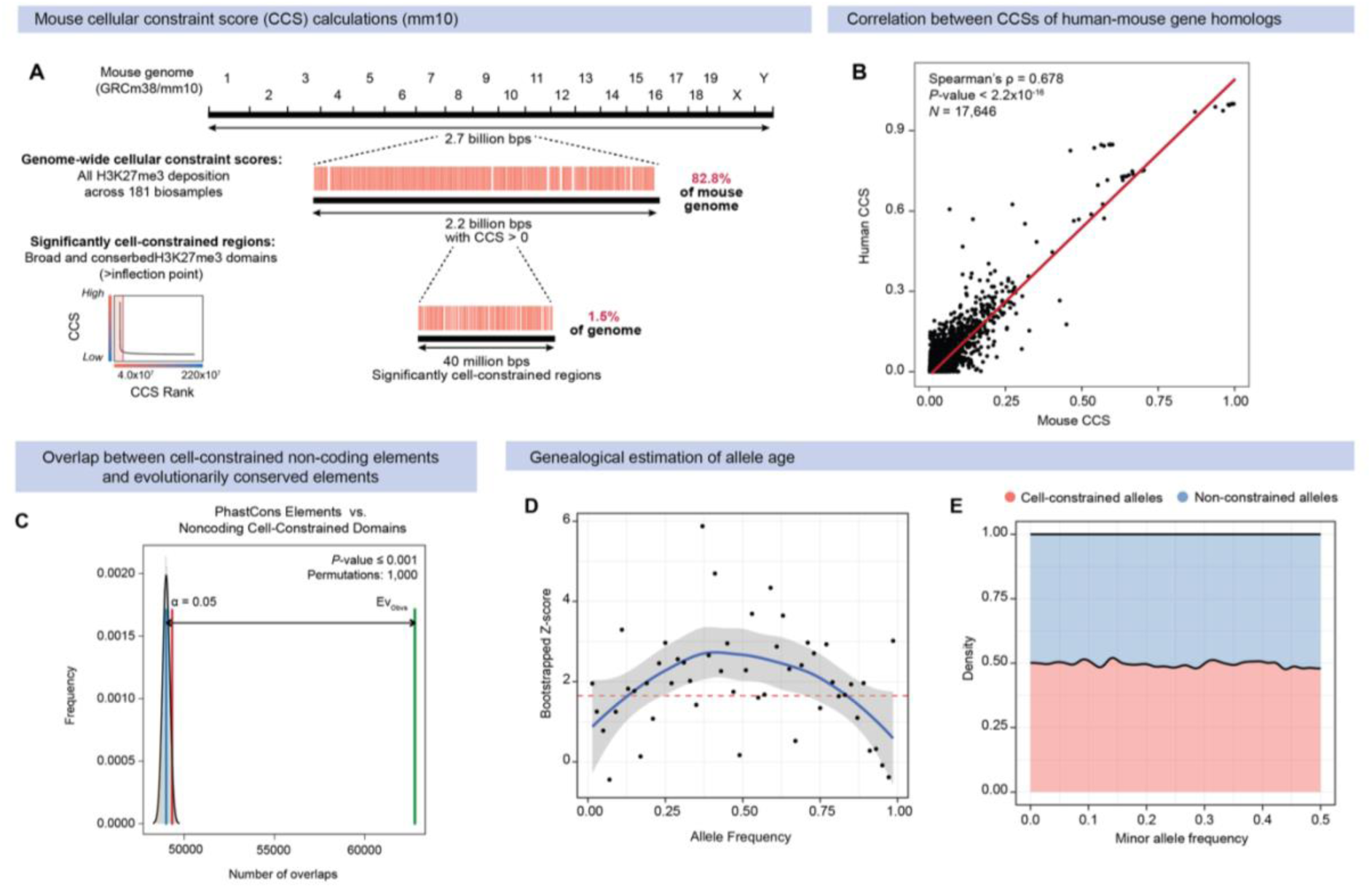
Mouse cellular constraint calculation and evolutionary analyses. **(A)** Single-base cellular constraint score calculation for the mouse genome (mm10). CCS reference metric covers 82.8% (2.2 billion bps) of the mouse genome, of which a subset of 1.5% (40 million bps) are defined as significantly cell-constrained bases. **(B)** Correlation between human and mouse cellular constraint calculated for homologous genes mapped between the human genome (hg19) and the mouse genome (mm10) (Spearman’s ρ = 0.678; *P*<2.2×10^-16^, Wilcoxon’s rank sum test). **(C)** Permutation tests showing enrichment of only non-coding cell-constrained domains in regions with PhastCons conserved elements (*P* ≤ 0.001) based on *n* = 1,000 random permutations. The average expected number of overlaps (blue) is compared against the observed number of overlaps (green), with α = 0.05 confidence interval indicated (red). **(D)** Bootstrapped Z-score indicating statistical significance at each MAF bin comparing allele age of significantly cell-constrained versus non-constrained variants. **(E)** Distribution of significantly cell-constrained and non-constrained variants across a minor allele frequency (MAF) range.

**Supplementary Figure 4.**
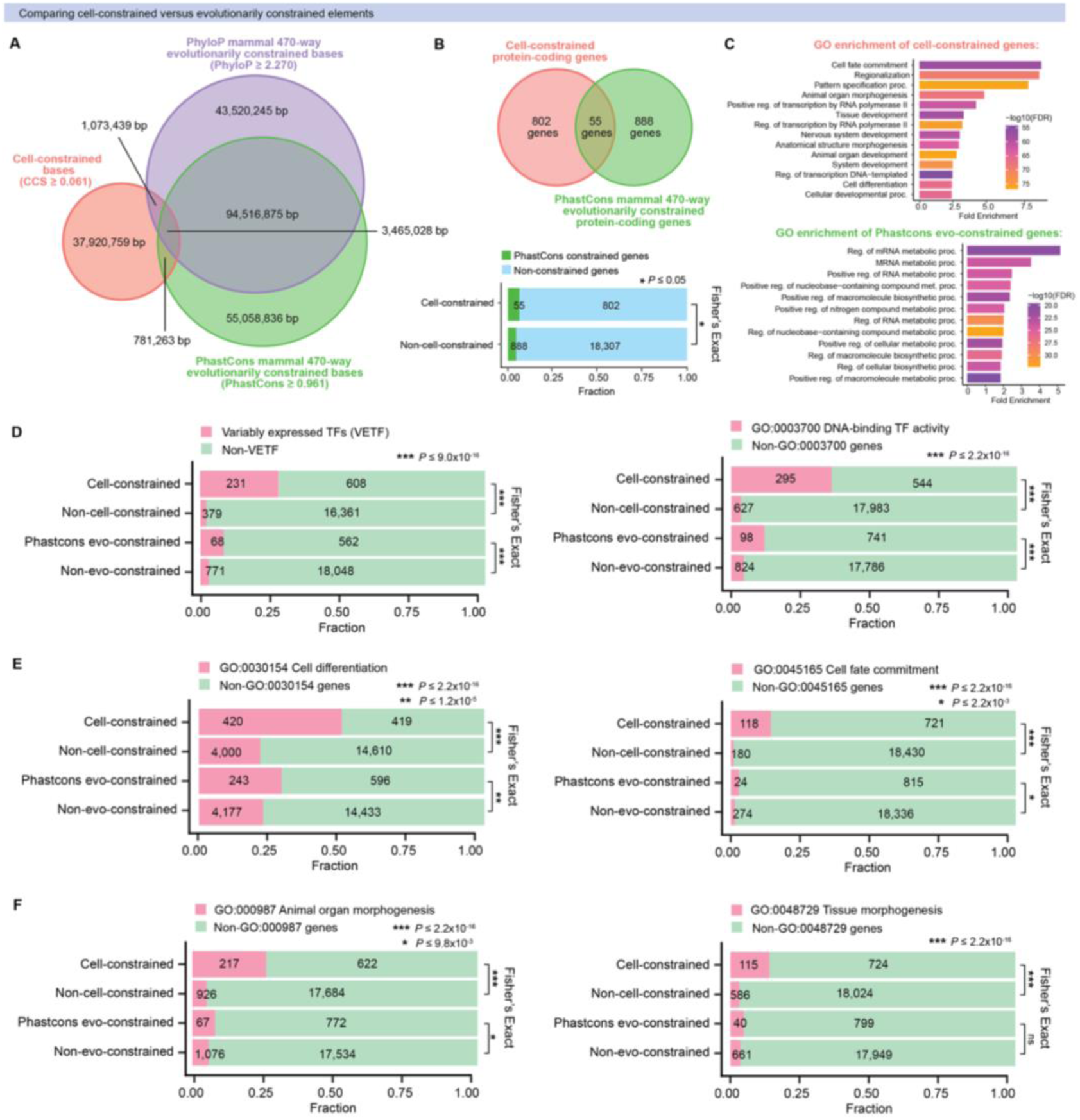
Comparison of cell-constrained versus evolutionarily constrained genes. **(A)** Euler diagram of overlap between genome-wide cell-constrained bases (CCS ≥ 0.061), PhastCons mammal 470-way evolutionarily constrained bases (PhastCons ≥ 0.961) and PhyloP mammal 470-way evolutionarily constrained bases (PhyloP ≥ 2.270). (**B**) Overlap between cell-constrained protein-coding genes (*N **=*** 857) and PhastCons mammal 470-way evolutionarily constrained protein-coding genes (*N **=*** 943). (**C**) Gene ontology (GO) enrichment of biological processes associated with the top 100 cell-constrained protein-coding genes versus PhastCons mammal 470-way evolutionarily constrained protein-coding genes. (**D-F**) Comparing the enrichment of representative gene-sets using cellular constraint versus PhastCons mammal 470-way evolutionarily constraint (\*\*\**P≤*2.2×10^-16^, two-tailed Fisher’s exact test).

**Supplementary Figure 5.**
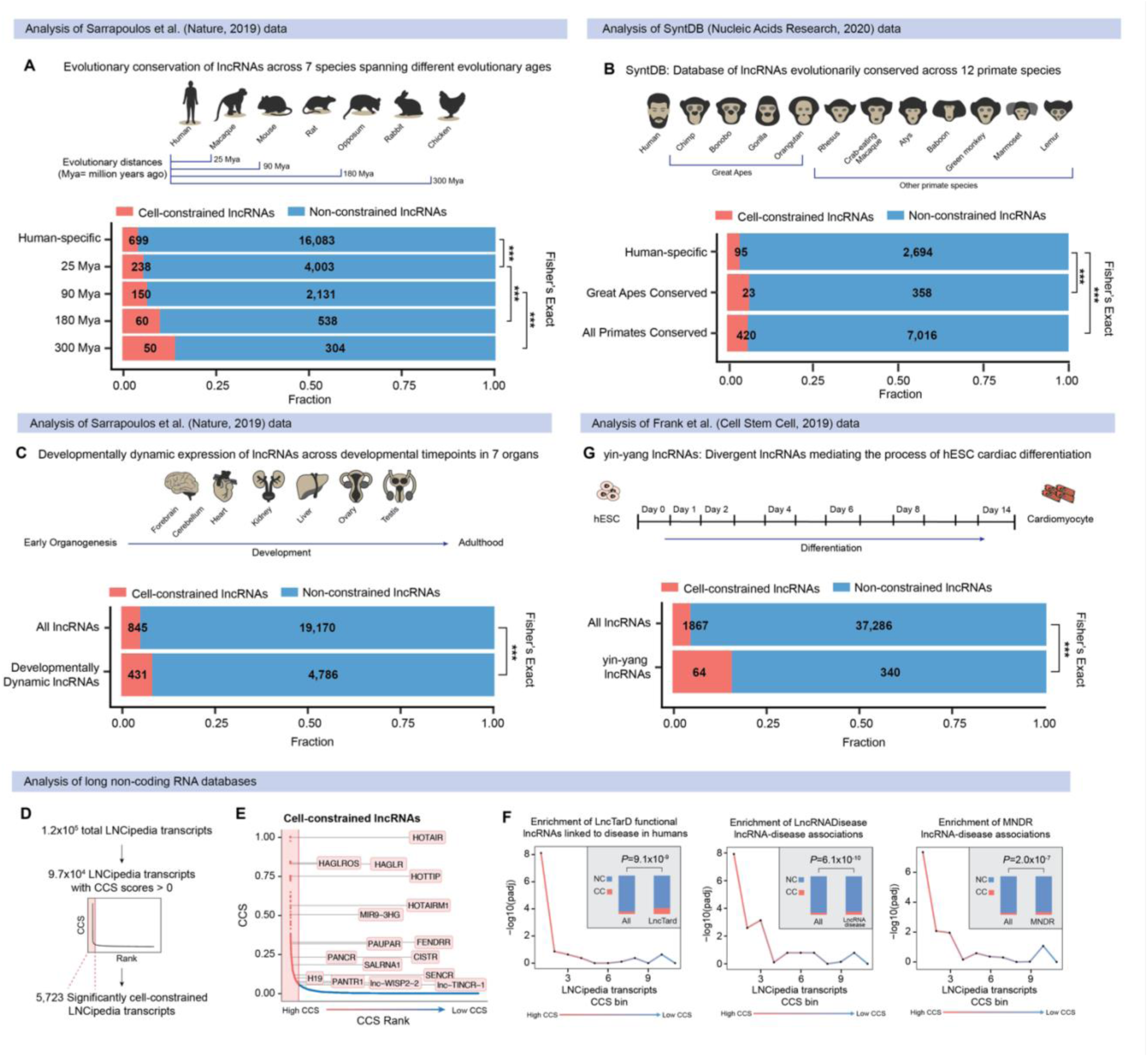
Enrichment of functional lncRNAs regulating development and disease. (**A)** Fraction of cell-constrained long non-coding RNAs (lncRNAs) represented in human lncRNAs conserved at different evolutionary ages from Sarrapoulos *et al.* (2019) ^30^ (\*\*\**P*<0.001, two-tailed Fisher’s exact test). (**B**) Fraction of cell-constrained lncRNAs represented in human lncRNAs conserved across 12 primate species from the SyntDB database ^53^ (\*\*\**P*<0.001, two-tailed Fisher’s exact test). (**C)** Fraction of cell-constrained lncRNAs represented in developmentally dynamic lncRNAs from Sarrapoulos *et al.* (2019) ^30^ (*P=*4.63×10^-29^, two-tailed Fisher’s exact test). **(D)** Cellular constraint score (CCS) generation for all LNCipedia ^92^ transcripts and calculation of inflection point to define a set of significantly cell-constrained lncRNAs (*n* = 5,723). **(E)** Cellular constraint-based prioritization of lncRNAs with well-known roles in regulating diverse developmental and disease processes. **(F)** Cellular constraint enriches for lncRNAs associated with disease from the LncTarD ^61^, LncRNADisease ^86^, and MNDR^87^ databases. Enrichment is plotted for lncRNAs sorted and grouped into percentile bins based on CCS metric (one-tailed Fisher’s exact test). Inset represents fraction of cell-constrained lncRNAs for each database (\*\*\**P*<0.001, two-tailed Fisher’s exact test). **(G)** Fraction of cell-constrained lncRNAs represented in yin-yang divergent lncRNAs from Frank *et al.* (2019) ^93^ (*P=*9.86×10^-17^, two-tailed Fisher’s exact test).

**Supplementary Figure 6:**
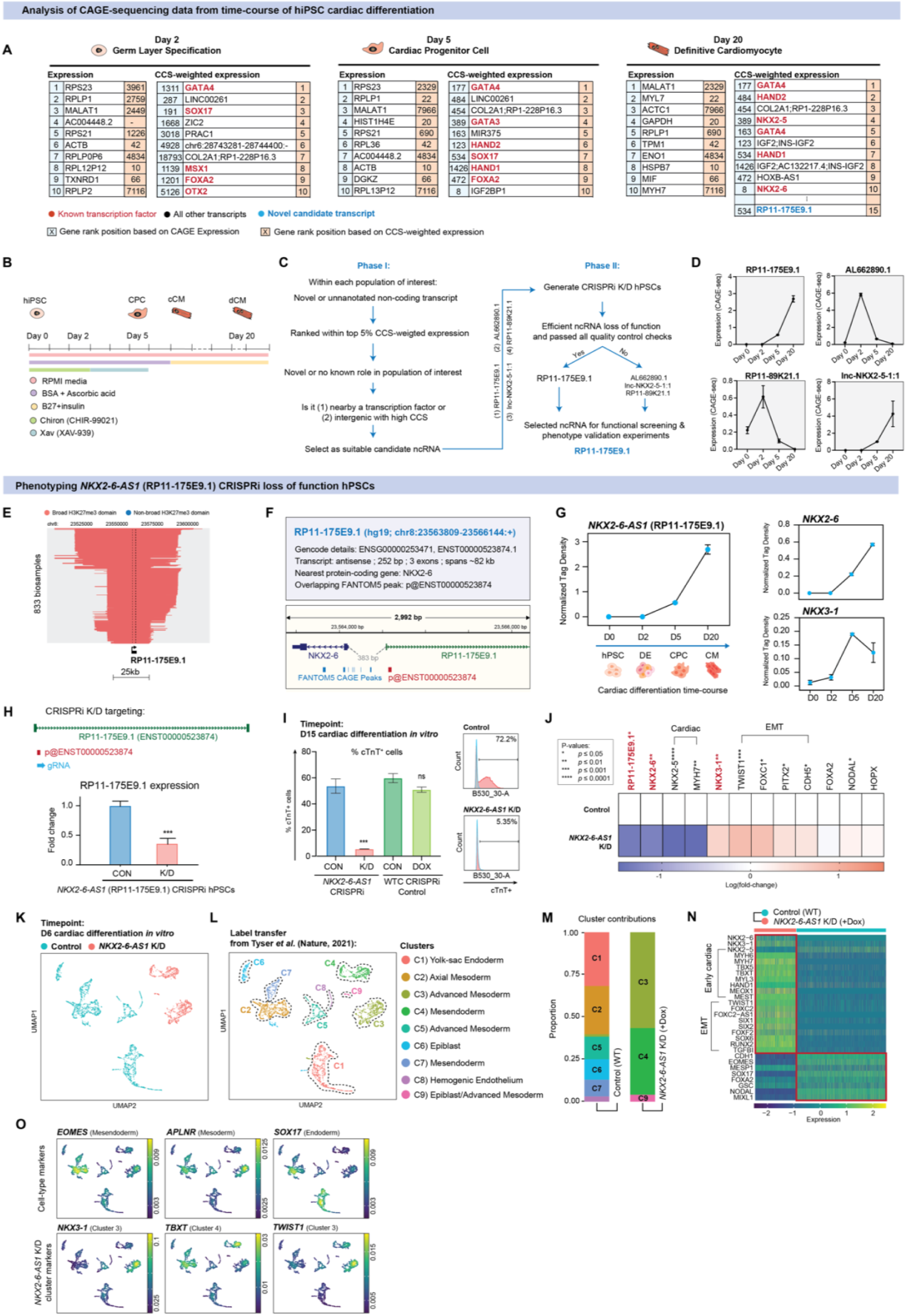
Experimental validation of cell-constrained divergent lncRNA *NKX2-6-AS1* during human cardiac differentiation. **(A)** Analysis of CAGE-seq data from time-course of hiPSC cardiac differentiation spanning pluripotency (Day 0), germ layer (Day 2), cardiac progenitors (Day 5) to definitive cardiomyocyte (Day 20) specification. Tables represent top 10 CAGE-defined Transcription Start Sites (TSSs) prioritized by rank of expression or cellular constraint-weighted expression. **(B)** Schematic of hiPSC cardiac differentiation protocol and relevant time-points. hiPSC = human induced pluripotent stem cell; CPC = cardiac progenitor cells; cCM = committed cardiomyocyte; dCM = definitive cardiomyocyte. **(C)** Pipeline for novel lncRNA candidate selection. **(D)** Time-course of candidate lncRNA transcript expression (normalized tag density) across different timepoint during hiPSC cardiac differentiation. Shown are the candidate lncRNAs *RP11-175E9.1, AL662890.1, lnc-NKX2-5-1:1, RP11-89K21.2* selected for CRISPRi K/D line generation. **(E)** Aligned ChIP-seq data from 833 EpiMap biosamples, showing H3K27me3 deposition over *RP11-175E9.1* stacked vertically and sorted on breadth. Domains ranked in the top 5% within each biosample are considered broad (red). **(F)** Genome browser representation of *NKX2-6-AS1* locus (*RP11-175E9.1*; chr8:23563809-23566144; hg19). **(G)** Comparative expression (normalized CAGE tag density) of *NKX2-6-AS1* (left), *NKX2-6* (top right) and *NKX3-1* (bottom right) across *in vitro* time-points. Data are represented as mean ± SEM. **(H)** CRISPRi knockdown (K/D) targeting of p1@ENST00000523874 to generate *NKX2-6-AS1* loss-of-function hPSCs (top). qPCR analysis of *RP11-175E9.1* transcript abundance in Control (WT) versus *NKX2-6-AS1* CRISPRi K/D hPSCs (*n* = 3 biological and *n* = 2 technical replicates) (bottom). **(I)** Day 15 FACs analysis of cTnT+ cardiomyocytes derived from Control versus *NKX2-6-AS1* K/D cells (*n* = 3 biological and *n* = 2 technical replicates; *P*=0.0003). Control and doxycycline-(dox-) treated WT cells are also represented to account for dox-effect (*n* = 3 biological and *n* = 2 technical replicates; ns) (**J**) qPCR analysis of panel of gene markers in Control versus *NKX2-6-AS1* K/D cells (*n* = 5-6 biological and *n* = 2 technical replicates). **(K)** Single-cell RNA-seq (scRNA-seq) UMAP of 3,333 cells profiling Control versus *NKX2-6-AS1* K/D conditions at Day 6 of cardiac differentiation *in vitro*. Cells are coloured by experimental condition. (**L**) Single-cell UMAP representation of 9 major subpopulations and their assigned cell-type identities based on label transfer from Tyser *et al*.^104^. Cells are coloured by predicted clusters. **(M)** Barplot representing the proportion of cells within Control or *NKX2-6-AS1* K/D conditions contributing to the overall diversity of subpopulations derived. **(N)** Heatmap of selected differentially expressed genes comparing Control versus *NKX2-6-AS1* K/D cells. Data represents z-score normalized counts scaled by row. **(O)** Density plot showing the expression of cluster biomarkers *EOMES, APLNR, SOX17, NKX3-1, TBXT* and *TWIST1*. Colours refer to gene-weighted kernel density as estimated by using R package Nebulosa.

