## Supplementary Material for "Epigenetic constraint of cellular genomes evolutionarily links genetic variation to function"

Enakshi Sinniah *et al*.

#### List of Supplementary Figures and Tables

**Supplementary Figure 1:** Cross-cell-type histone modification ChIP-seq alignment across diverse cell- and tissue-types.

**Supplementary Figure 2:** General characterization of human cellular constraint metric.

**Supplementary Figure 3:** Mouse cellular constraint calculation and evolutionary analyses.

**Supplementary Figure 4:** Comparison of cell-constrained versus evolutionarily constrained genes.

**Supplementary Figure 5:** Enrichment of functional lncRNAs regulating development and disease.

**Supplementary Figure 6:** Experimental validation of cell-constrained divergent lncRNA *NKX2-6-AS1* during human cardiac differentiation.

**Supplementary Table 1.** Human cellular constraint scores for Refseq protein-coding genes.

**Supplementary Table 2.** Human and mouse cellular constraint scores for homologous genes mapped between species.

**Supplementary Table 3.** Human cellular constraint scores for LNCipedia transcripts.

**Supplementary Table 4.** Cellular constraint weighted abundance of CAGE-seq transcripts from time-course of hiPSC cardiac differentiation.

**Supplementary Table 5.** Significantly cell-constrained GTEx cis-eQTLs across 49 cell- and tissue-types.

*See separate .xlsx or .txt files attached for the supplementary tables.*

#### Supplementary Text

*Supplementary text and references accompanying the manuscript are provided at the end of this document.*

#### Supplementary Figures

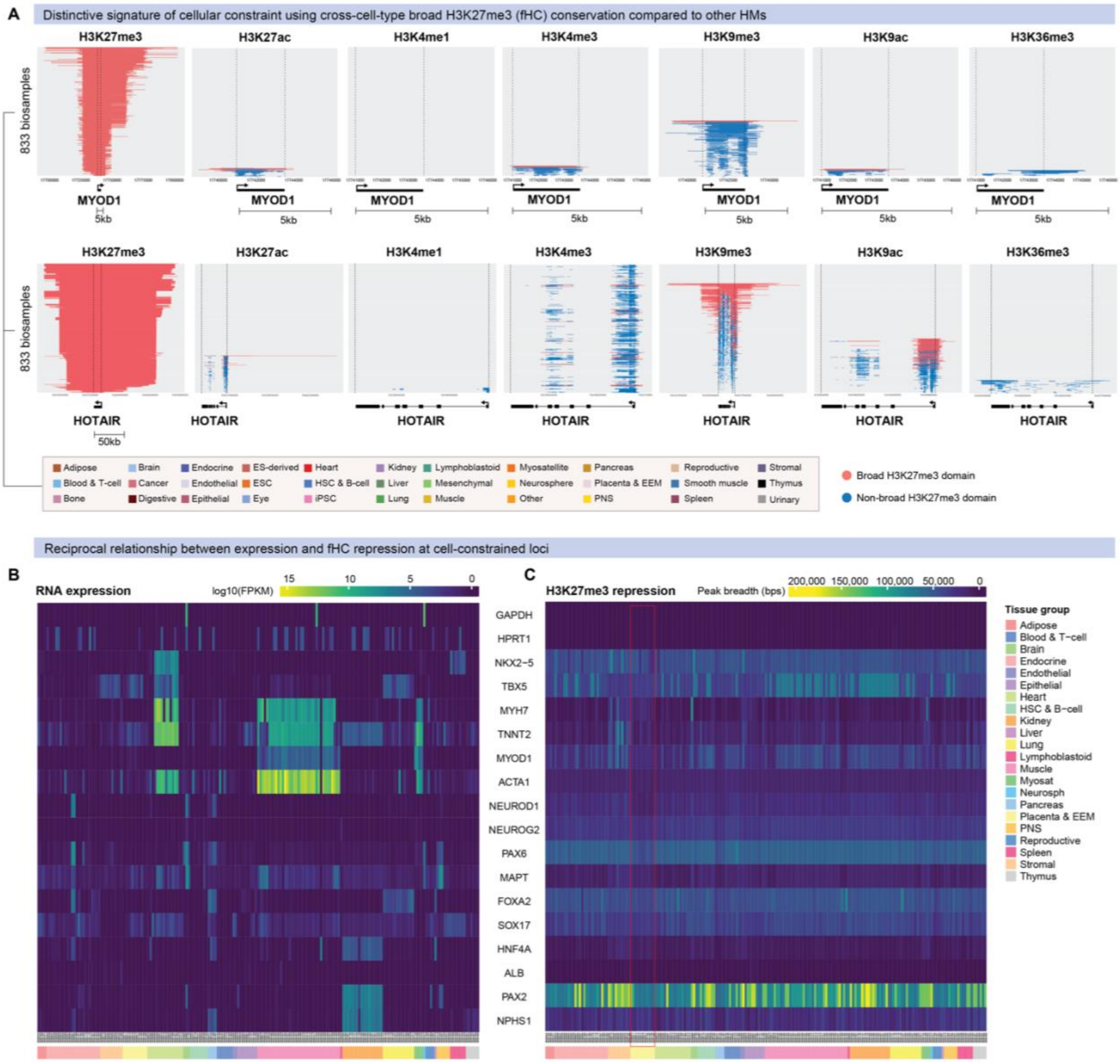


**Supplementary Figure 1. Cross-cell-type histone modification ChIP-seq alignment across diverse cell- and tissue-types.**
**(A)** Plots display aligned ChIP-seq peaks from 833 EpiMap biosamples, showing deposition of seven HMs over a locus stacked vertically and sorted on breadth. Domains ranked in the top 5% within each biosample are considered broad (red) and the remainder non-broad (blue). **(B)** Heatmaps displaying RNA expression (log10 FPKM) (left) and H3K27me3 repression (peak breadth) (right) of representative genes.

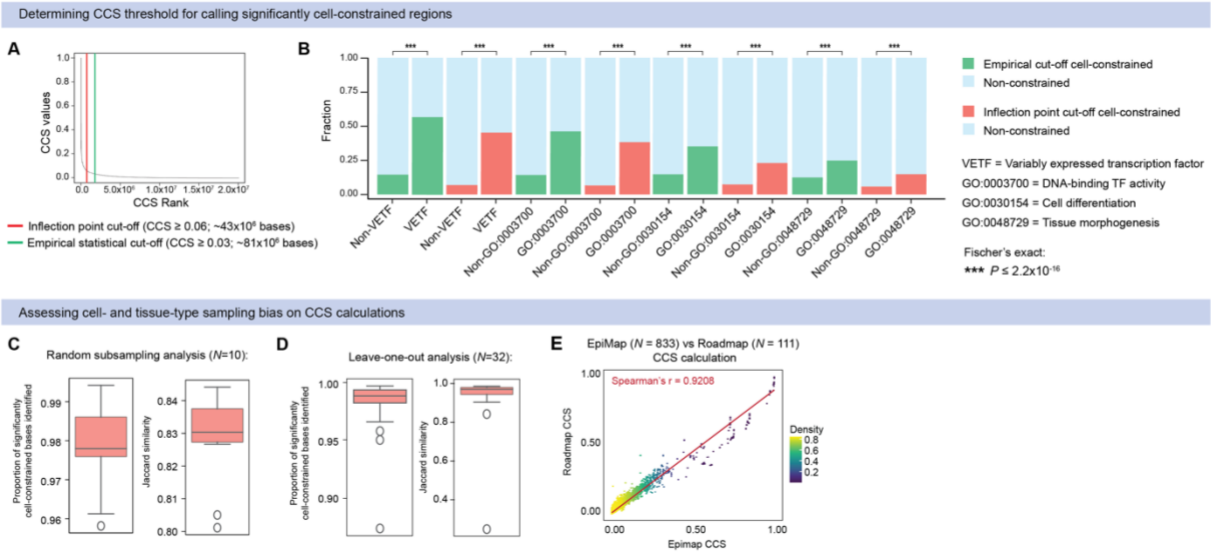


**Supplementary Figure 2. General characterization of human cellular constraint metric.** **(A)** Distribution of single-base CCS values showing thresholds for calling significantly cell-constrained bases using the inflection point cut-off (CCS ≥ 0.06; ~43x$10^6$ bases) (red line) vs. an empirical statistical cut-off (CCS ≥ 0.03, ~81x$10^6$ bases) (green line). (**B**) Comparing the enrichment of representative gene-sets using the inflection point versus an empirical statistical cut-off to determine significantly cell-constrained bases (***$P \leq 2.2 \times 10^{-16}$, two-tailed Fisher's exact test). (**C-E**) Evaluating cell- or tissue-type bias within biosamples effect on CCS calculations. (**C**) Bootstrapping analysis via random resampling of 833 biosamples (*N*= 10 permutations) showing proportion (left) and jaccard similarity index (right) of significantly cell-constrained bases identified. (**D**) Leave-one-out analysis via iterative removal of all biosamples from each of the 33 tissue-groups represented (*N*= 32 permutations) showing proportion (left) and jaccard similarity index (right) of significantly cell-constrained bases identified. (**E**) Correlation between cellular constraint scores calculated used epigenomes from EpiMap (*N*= 833 biosamples) versus Roadmap (*N*= 111 biosamples) (Spearman's *r*= 0.9208, *p*=<0.0001).

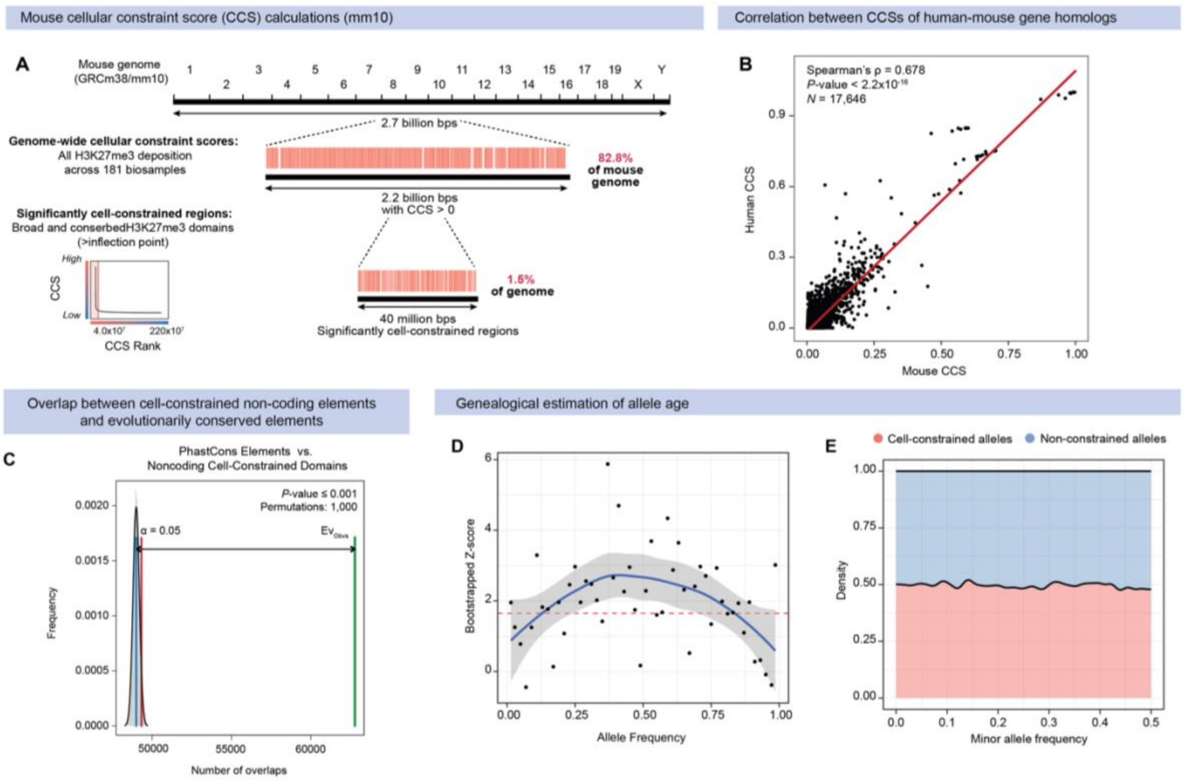


**Supplementary Figure 3. Mouse cellular constraint calculation and evolutionary analyses.**

**(A)** Single-base cellular constraint score calculation for the mouse genome (mm10). CCS reference metric covers 82.8% (2.2 billion bps) of the mouse genome, of which a subset of 1.5% (40 million bps) are defined as significantly cell-constrained bases. **(B)** Correlation between human and mouse cellular constraint calculated for homologous genes mapped between the human genome (hg19) and the mouse genome (mm10) (Spearman's ρ = 0.678; $P<2.2x10^{-16}$, Wilcoxon's rank sum test). **(C)** Permutation tests showing enrichment of only non-coding cell-constrained domains in regions with PhastCons conserved elements ($P \leq 0.001$) based on $n$ = 1,000 random permutations. The average expected number of overlaps (blue) is compared against the observed number of overlaps (green), with α = 0.05 confidence interval indicated (red). **(D)** Bootstrapped Z-score indicating statistical significance at each MAF bin comparing allele age of significantly cell-constrained versus non-constrained variants. **(E)** Distribution of significantly cell-constrained and non-constrained variants across a minor allele frequency (MAF) range.

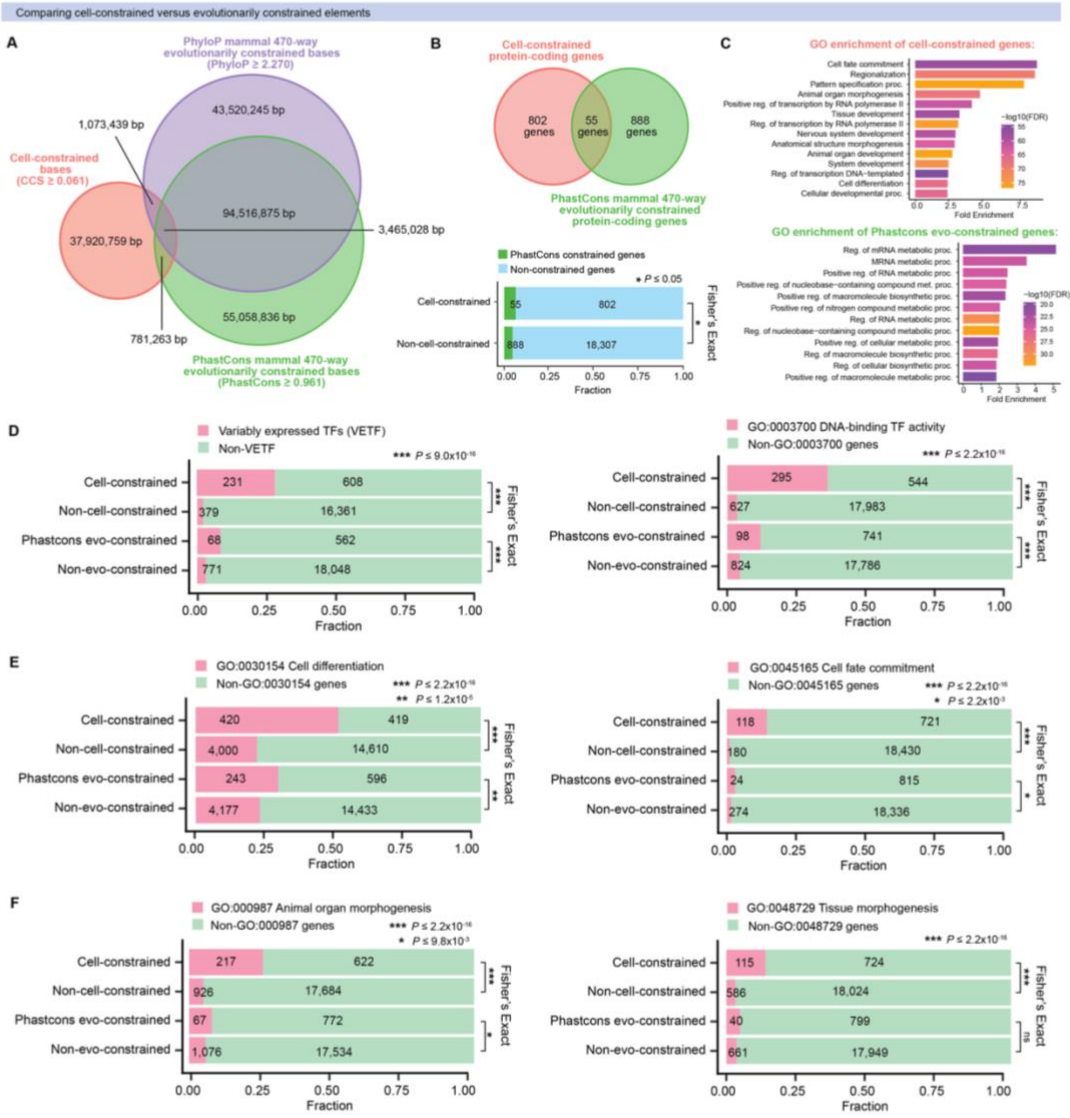


**Supplementary Figure 4. Comparison of cell-constrained versus evolutionarily constrained genes.**

**(A)** Euler diagram of overlap between genome-wide cell-constrained bases (CCS ≥ 0.061), PhastCons mammal 470-way evolutionarily constrained bases (PhastCons ≥ 0.961) and PhyloP mammal 470-way evolutionarily constrained bases (PhyloP ≥ 2.270). (**B**) Overlap between cell-constrained protein-coding genes ($N$ = 857) and PhastCons mammal 470-way evolutionarily constrained protein-coding genes ($N$ = 943). (**C**) Gene ontology (GO) enrichment of biological processes associated with the top 100 cell-constrained protein-coding genes versus PhastCons mammal 470-way evolutionarily constrained protein-coding genes. (**D-F**) Comparing the enrichment of representative gene-sets using cellular constraint versus

PhastCons mammal 470-way evolutionarily constraint (****$P \leq 2.2 \times 10^{-16}$, two-tailed Fisher's exact test).

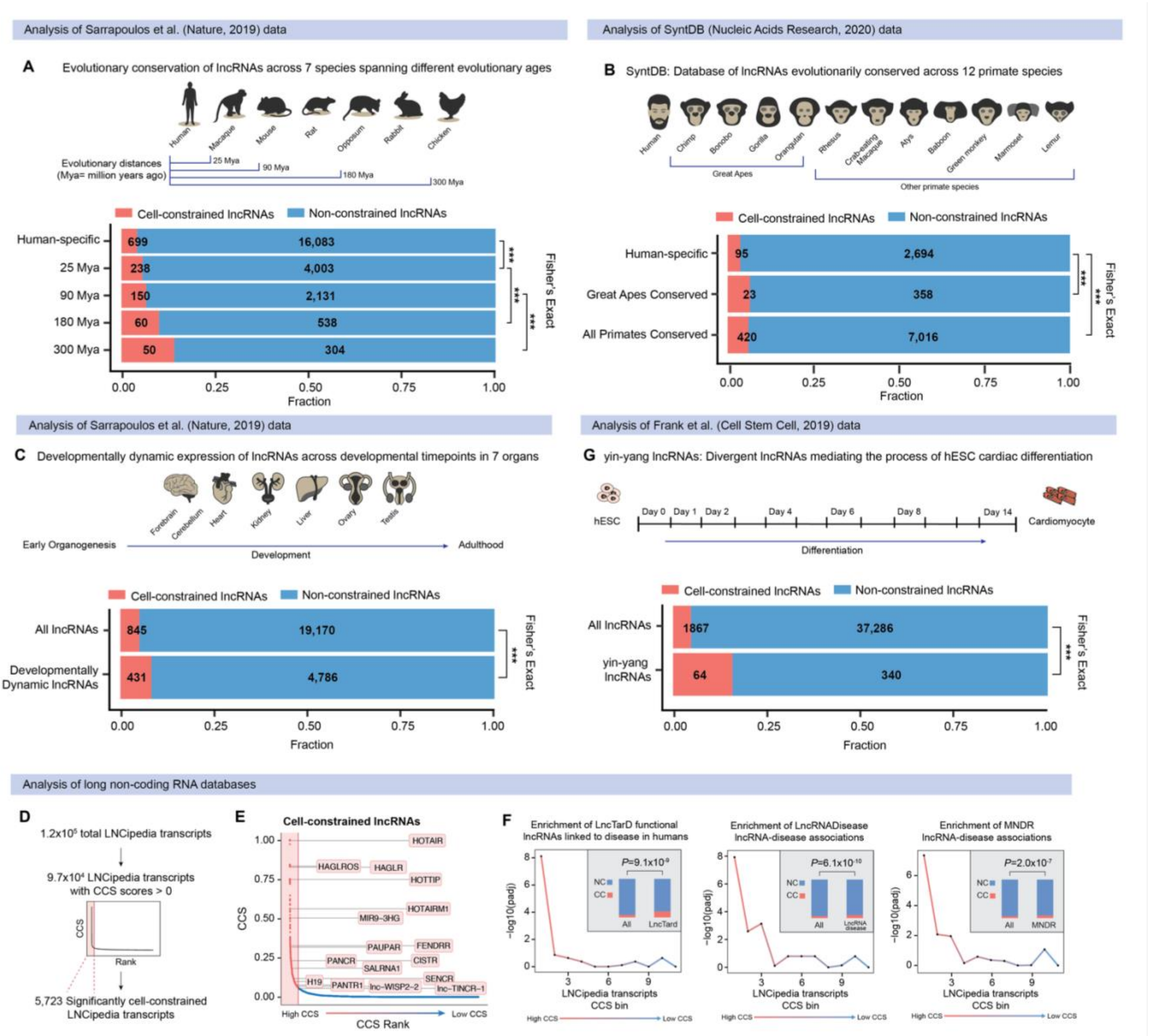


**Supplementary Figure 5. Enrichment of functional lncRNAs regulating development and disease.**

**(A)** Fraction of cell-constrained long non-coding RNAs (lncRNAs) represented in human lncRNAs conserved at different evolutionary ages from Sarrapoulos *et al.* (2019) [1] (***$P$<0.001, two-tailed Fisher's exact test). (**B**) Fraction of cell-constrained lncRNAs represented in human lncRNAs conserved across 12 primate species from the SyntDB database [2] (***$P$<0.001, two-tailed Fisher's exact test). **(C)** Fraction of cell-constrained lncRNAs represented in developmentally dynamic lncRNAs from Sarrapoulos *et al.* (2019) [1] ($P$=4.63x$10^{-29}$, two-tailed Fisher's exact test). **(D)** Cellular constraint score (CCS) generation for all LNCipedia [3] transcripts and calculation of inflection point to define a set of significantly cell-constrained lncRNAs ($n$ = 5,723). **(E)** Cellular constraint-based prioritization of lncRNAs with well-known roles in regulating diverse developmental and disease processes. **(F)** Cellular constraint enriches for lncRNAs associated with disease from the LncTarD [4], LncRNADisease [5], and MNDR[6] databases. Enrichment is plotted for lncRNAs sorted and grouped into percentile bins based on CCS metric (one-tailed Fisher's exact test). Inset represents fraction of cell-constrained lncRNAs for each database (***$P$<0.001, two-tailed Fisher's exact test). **(G)** Fraction of cell-constrained lncRNAs represented in yin-yang divergent lncRNAs from Frank *et al.* (2019) [7] ($P$=9.86x10$^{-17}$, two-tailed Fisher's exact test).

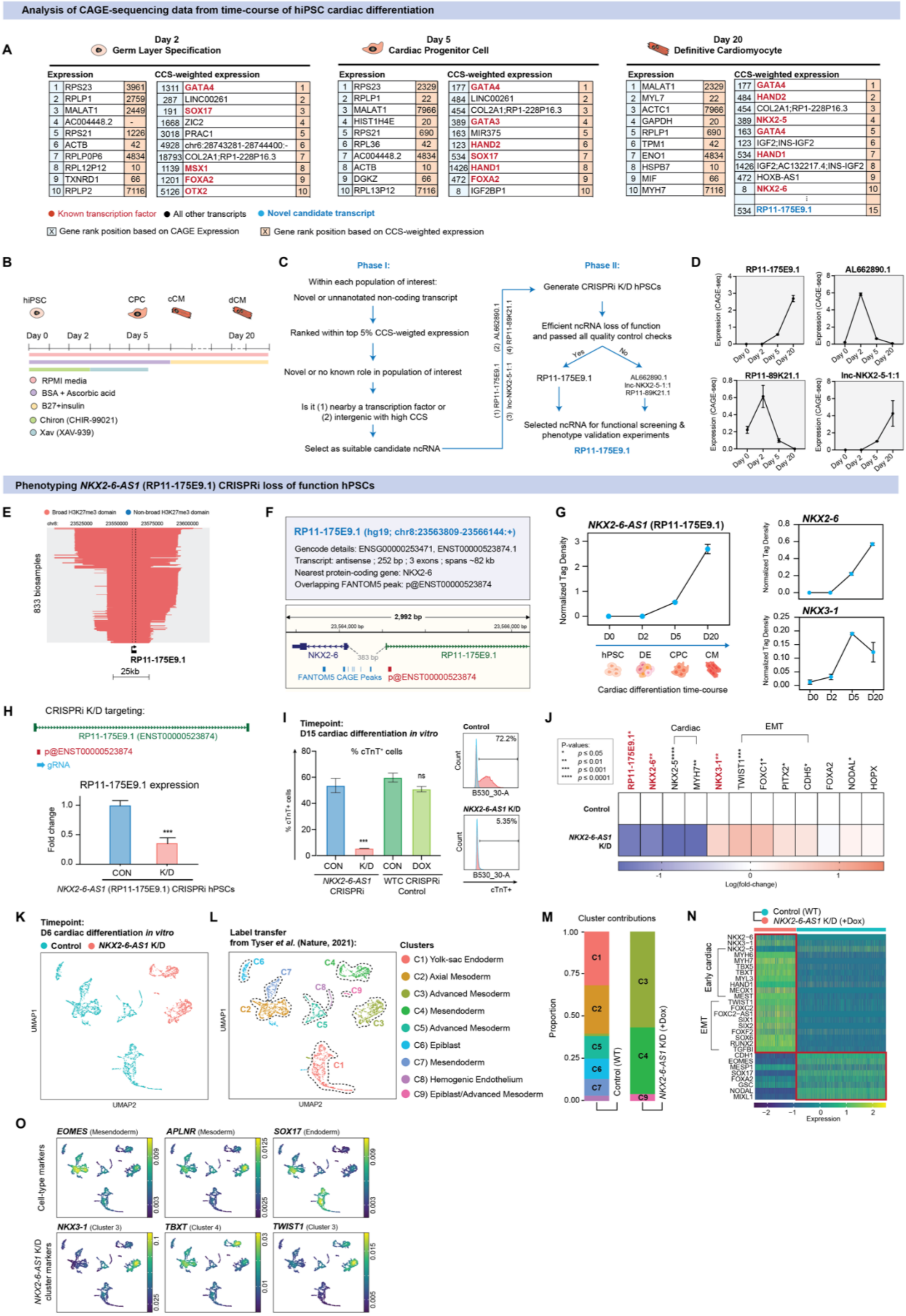

Analysis of CAGE-sequencing data from time-course of hiPSC cardiac differentiation
A
Day 2 Germ Layer Specification
Day 5 Cardiac Progenitor Cell
Day 20 Definitive Cardiomyocyte
Expression
CCS-weighted expression
1 RPS23 3961 | 1311 GATA4 1
2 RPLP1 2759 | 287 LINC00261 2
3 MALAT1 2449 | 191 SOX17 3
4 AC004448.2 - | 1668 ZIC2 4
5 RPS21 1226 | 3018 PRAC1 5
6 ACTB 42 | 4928 chr6:28743281-28744400:- 6
7 RPLP0P6 4834 | 18793 COL2A1;RP1-228P16.3 7
8 RPL12P12 10 | 1139 MSX1 8
9 TXNRD1 66 | 1201 FOXA2 9
10 RPLP2 7116 | 5126 OTX2 10
1 RPS23 2329 | 177 GATA4 1
2 RPLP1 22 | 484 LINC00261 2
3 MALAT1 7966 | 454 COL2A1;RP1-228P16.3 3
4 HIST1H4E 20 | 389 GATA3 4
5 RPS21 690 | 163 MIR375 5
6 RPL36 42 | 123 HAND2 6
7 AC004448.2 4834 | 534 SOX17 7
8 ACTB 10 | 1426 HAND1 8
9 DGKZ 66 | 472 FOXA2 9
10 RPL13P12 7116 | 8 IGF2BP1 10
1 MALAT1 2329 | 177 GATA4 1
2 MYL7 22 | 484 HAND2 2
3 ACTC1 7966 | 454 COL2A1;RP1-228P16.3 3
4 GAPDH 20 | 389 NKX2-5 4
5 RPLP1 690 | 163 GATA4 5
6 TPM1 42 | 123 IGF2;INS-IGF2 6
7 ENO1 4834 | 534 HAND1 7
8 HSPB7 10 | 1426 IGF2;AC132217.4;INS-IGF2 8
9 MIF 66 | 472 HOXB-AS1 9
10 MYH7 7116 | 8 NKX2-6 10
534 RP11-175E9.1 15
● Known transcription factor ● All other transcripts ● Novel candidate transcript
X Gene rank position based on CAGE Expression
X Gene rank position based on CCS-weighted expression
B
hiPSC CPC cCM dCM
Day 0 Day 2 Day 5 Day 20
RPMI media
BSA + Ascorbic acid
B27+insulin
Chiron (CHIR-99021)
Xav (XAV-939)
C
Phase I:
Within each population of interest:
Novel or unannotated non-coding transcript
Ranked within top 5% CCS-weigted expression
Novel or no known role in population of interest
Is it (1) nearby a transcription factor or (2) intergenic with high CCS
Select as suitable candidate ncRNA
(1) RP11-175E9.1 (2) AL662890.1 (3) lnc-NKX2-5-1:1 (4) RP11-89K21.1
Phase II:
Generate CRISPRi K/D hPSCs
Efficient ncRNA loss of function and passed all quality control checks
Yes No
RP11-175E9.1
AL662890.1 lnc-NKX2-5-1:1 RP11-89K21.1
Selected ncRNA for functional screening & phenotype validation experiments
RP11-175E9.1
D
RP11-175E9.1
AL662890.1
RP11-89K21.1
lnc-NKX2-5-1:1
Expression (CAGE-seq)
Day 0 Day 2 Day 5 Day 20
Phenotyping NKX2-6-AS1 (RP11-175E9.1) CRISPRi loss of function hPSCs
E
Broad H3K27me3 domain
Non-broad H3K27me3 domain
833 biosamples
RP11-175E9.1
25kb
F
RP11-175E9.1 (hg19; chr8:23563809-23566144:+)
Gencode details: ENSG00000253471, ENST00000523874.1
Transcript: antisense ; 252 bp ; 3 exons ; spans ~82 kb
Nearest protein-coding gene: NKX2-6
Overlapping FANTOM5 peak: p@ENST00000523874
2,992 bp
NKX2-6
383 bp
RP11-175E9.1
FANTOM5 CAGE Peaks
p@ENST00000523874
G
NKX2-6-AS1 (RP11-175E9.1)
Normalized Tag Density
D0 D2 D5 D20
hPSC DE CPC CM
Cardiac differentiation time-course
NKX2-6
NKX3-1
H
CRISPRi K/D targeting:
RP11-175E9.1 (ENST00000523874)
p@ENST00000523874
gRNA
RP11-175E9.1 expression
Fold change
CON K/D
NKX2-6-AS1 (RP11-175E9.1) CRISPRi hPSCs
I
Timepoint:
D15 cardiac differentiation in vitro
% cTnT+ cells
CON K/D CON DOX
NKX2-6-AS1 CRISPRi
WTC CRISPRi Control
Control 72.2%
NKX2-6-AS1 K/D 5.35%
B530_30-A
cTnT+
J
P-values:
* p ≤ 0.05
** p ≤ 0.01
*** p ≤ 0.001
**** p ≤ 0.0001
Cardiac
EMT
RP11-175E9.1* NKX2-6** NKX2-5**** MYH7** NKX3-1** TWIST1*** FOXC1* PITX2* CDH5* FOXA2 NODAL* HOPX
Control
NKX2-6-AS1 K/D
Log(fold-change)
K
Timepoint:
D6 cardiac differentiation in vitro
Control NKX2-6-AS1 K/D
UMAP1
UMAP2
L
Label transfer from Tyser et al. (Nature, 2021):
Clusters
C1) Yolk-sac Endoderm
C2) Axial Mesoderm
C3) Advanced Mesoderm
C4) Mesendoderm
C5) Advanced Mesoderm
C6) Epiblast
C7) Mesendoderm
C8) Hemogenic Endothelium
C9) Epiblast/Advanced Mesoderm
M
Cluster contributions
Proportion
Control (WT)
NKX2-6-AS1 K/D (+Dox)
N
Control (WT)
NKX2-6-AS1 K/D (+Dox)
Early cardiac
EMT
NKX2-6 NKX3-1 NKX2-5 MYH6 MYH7 TBX5 TBXT MYL3 HAND1 MEOX1 MEST TWIST1 FOXC2 FOXC2-AS1 SIX1 SIX2 FOXF2 SOX6 RUNX2 TGFBI CDH1 EOMES MESP1 SOX17 FOXA2 GSC NODAL MIXL1
Expression
O
EOMES (Mesendoderm)
APLNR (Mesoderm)
SOX17 (Endoderm)
NKX3-1 (Cluster 3)
TBXT (Cluster 4)
TWIST1 (Cluster 3)
Cell-type markers
NKX2-6-AS1 K/D cluster markers

### Supplementary Text

***Validation of cell-constrained lncRNA NKX2-6-AS1 predicted to regulate cardiac fate.***
(Figure 4G and S6)
In this section, we expand on the description of results from genetic loss-of-function perturbation experiments of *NKX2-6-AS1* (*RP11-175E9.1*) during *in vitro* hiPSC cardiac differentiation.

Divergent lncRNAs are transcribed from the bidirectional promoters of nearby protein-coding genes and have emerged as critical regulators of cellular differentiation [9,10]. Since broad and conserved H3K27me3 domains selectively repress developmental regulatory genes [11], we hypothesized that cell-constrained loci are likely to be enriched for divergent lncRNAs governing cell identity. In support of this, we observed significant enrichment of divergent yin-yang lncRNAs (yylncRNAs) [7] associated with regulating cardiac lineage specification *in vitro* (**Figure S5G,** $P$=9.86x10$^{-17}$, two-sided Fisher's exact test).

Using cellular constraint to weight transcript abundance at each differentiation timepoint (**Figure S6A** and **Supplementary Table 4**), we identified *RP11-175E9.1* (ENST00000523874.1) as a CCS-prioritized candidate novel divergent lncRNA transcript predicted to regulate cardiogenesis (**Figure** S6E). *RP11-175E9.1* is an antisense lncRNA transcribed ~383bp directly upstream of *NKX2-6* and nearby *NKX3-1*, a pair of TFs with known roles in regulating cardiac [12-14] and prostate development [15,16], respectively (**Figure S6F**). Using published promoter capture Hi-C (PCHi-C) data of human pluripotent stem cells (hPSCs) and hPSC-derived cardiomyocytes [17], we analyzed genome-wide chromatin interaction maps and found that *RP11-175E9.1* physically interacts with promoters of both *NKX3-1* and *NKX2-6,* suggesting it likely acts in *cis* to regulate the transcription of these nearby genes (**see Table in Methods**). *RP11-175E9.1* is first detected in the cardiac progenitor stage (D5) and is highest in late stages (D20) of cardiac commitment, displaying an expression pattern identical to that of *NKX2-6* (**Figure S6G**). On this basis, we considered *RP11-175E9.1* as a novel divergent lncRNA transcribed from the *NKX2-6* locus, which we consequently termed and refer to as *NKX2-6-AS1*.

For experimental perturbation of *NKX2-6-AS1*, we established CRISPRi loss-of-function (LOF) hPSCs for targeted knockdown of transcription at its CAGE-defined TSS (**Figure S6H**). First, we performed single cell transcriptional profiling of *NKX2-6-AS1* LOF and wild-type (WT) control hPSCs at early stages (day 6) of *in vitro* cardiac differentiation. After cell filtering and quality control, a total of 3,333 cells were captured, clustered, and projected in a two-dimensional space using Uniform Manifold Approximation and Projection (UMAP) for dimensionality reduction (**Figure S6K**). We annotated cell clusters using Seurat's label transfer function [18] based on annotations of the gastrulating human embryo from Tyser *et al.*[8] (**Figure S6L**). We found that *NKX2-6-AS1* knockdown (K/D) cells clustered independently, with little to no contribution to the endodermal cell populations derived from WT control cells (**Figure S6M**). Notably, we observed significant upregulation of *NKX3-1* expression in *NKX2-6-AS1* K/D cells, accompanied by the upregulation of markers of epithelial-to-mesenchymal transition (EMT) such as *TWIST1*, *FOXC2*, and *SIX1* (**Figures S6N & S6O**). *NKX3-1* is the earliest known marker of prostate epithelium development and is facilitated by EMT during embryogenesis [15,19,20], indicating that *NKX2-6-AS1* LOF at day 6 could promote prostate differentiation. Together, this data suggests that *NKX2-6-AS1* may act to suppress *NKX3-1* expression, to maintain cardiac fate upon divergent initiation of *NKX2-6* transcription during early stages of cardiac specification.

To further analyze the *NKX2-6-AS1* LOF phenotype, we focused on assessing its effect at late-stages (D15) of *in vitro* cardiac differentiation. We found that *NKX2-6-AS1* K/D hPSCs failed to differentiate into cardiomyocytes, with flow cytometry showing a significant depletion of cardiac troponin $T^+$ ($cTnT^+$) cells measured at day 15 as compared against dox-treated WT controls ($n$ = 3 biological and $n$ = 2 technical replicates each; $P$=0.0003) (**Figure S6I)**. As expected, we observed significant depletion of *NKX2-6* expression ($P\leq 0.005$) accompanied by downregulation of cardiac markers *NKX2-5* ($P\leq 0.0001$) and *MYH7* ($P$=0.0082) upon loss of *NKX2-6-AS1* (*RP11-175E9.1*) ($P$=0.0123) ($n$ = 5-6 biological and $n$ = 2 technical replicates each) (**Figure S6J**). Consistent with our scRNA-seq results, we observed significant enrichment of *NKX3-1* expression ($P$=0.0038) together with upregulation of prostate-related EMT markers *TWIST1* ($P$=0.0009), *FOXC1* ($P$=0.0203), *PITX2* ($P$=0.0214) and *CDH5* ($P$=0.0203) ($n$ = 5-6 biological and $n$ = 2 technical replicates each) (**Figure S6J)**. Together, this experimental validation evidences the functional role of *NKX2-6-AS1* as a divergent lncRNA regulator of cardiac lineage specification. Overall, these results demonstrate the utility of cellular constraint annotation to prioritize loci governing cell fate and discover novel non-coding regulators of cellular differentiation in a cell-type or cell-state specific manner.
